# Cep192 insufficiency underlies haploid instability in human cells

**DOI:** 10.64898/2026.03.18.712690

**Authors:** Koya Yoshizawa, Hemang Raj Singh, Paramasivam Kathirvel, Jin Zhu, Ryota Uehara

## Abstract

Mammalian somatic haploid cells offer advantages for genome engineering, yet rapid diploidization limits their utility. Here, we reveal that a haploidy-specific attenuation of mitotic spindle bipolarization, independent of previously characterized centrosome loss, underlies haploid instability in human cells. Comparative imaging and structure-function analyses demonstrate that the halved absolute dosage of the pericentriolar scaffolding protein Cep192 prevents its centrosomal accumulation to the threshold required for Aurora A–Eg5 axis. Consequently, haploids exhibit innate fragility in centrosome separation and spindle maintenance. Supplementing Cep192 restored spindle bipolarization to diploid levels and, when combined with genetic enhancement of the acentrosomal spindle pathway, profoundly stabilized the haploid state. Moreover, a genome-wide CRISPR-activation screen leveraging the above principle identified novel haploid-stabilizing genes, including the glutamate transporter SLC1A2. Our findings uncover an absolute-dosage scaling limit of mitotic scaffolding in haploids and establish genetic enhancement of spindle fidelity as an effective strategy for engineering stable animal haploid bioresources.

**Teaser:** Molecular elucidation of haploidy-linked fragility in mitotic spindle architecture enables engineering stable human haploid cells.

## Introduction

Haploid cells, which possess only a single copy of the genome, offer efficient genetic modification and direct genotype-phenotype correlation by eliminating interference from counter alleles (*1, 2*). This singular advantage has established haploid cells as invaluable bioresources for complex, multi-step genome editing and high-throughput genetic screens (*3–7*). While naturally occurring haploidy is prevalent in yeasts and plants (*8*), vertebrate somatic haploid cells generally arise only through irregular biological processes, such as parthenogenesis or drastic chromosome loss during carcinogenesis (*9*). Consequently, engineered mammalian somatic haploids hold immense promise (*9, 10*). However, animal somatic haploids universally exhibit marked chromosome instability, caused by mitotic chromosome missegregation, and poor viability compared to their diploid counterparts (*11–15*). This innate instability drives spontaneous diploidization, leading to the rapid and complete replacement of haploid populations by diploidized cells during routine culture (*11, 13*). Overcoming this biological barrier remains a major bottleneck in haploid cell technology (*16*).

Various treatments have been reported to improve the stability of animal haploid cells. Deletion of p53 or alleviation of endoplasmic reticulum stress improves haploid stability by suppressing haploidy-linked cell death, thereby slowing the replacement of haploid populations by diploidized populations (*11, 17–19*). However, to fundamentally resolve haploid instability, it is imperative to identify and correct the mechanical root causes of the chromosome missegregation that drives spontaneous diploidization. Previous studies indicated that haploidy-linked mitotic failure is driven, at least in part, by chronic, haploidy-specific centrosome loss (*13, 20*). While pharmacological extension of the S-phase can temporarily alleviate this loss and improve haploid retention, the utility of such approaches is severely hampered by drug toxicity and low efficiency (*20*). To engineer durable, versatile haploid cell resources, we must delineate the precise molecular mechanisms underpinning these mitotic defects and identify genetic targets to permanently restore mitotic fidelity.

Precise chromosome segregation relies on the dynamic assembly of a bipolar mitotic spindle. During mitotic entry, the pericentriolar material (PCM) expands, a process referred to as centrosome maturation, driven by the accumulation of key scaffolding proteins, including PCNT/Pericentrin, Cep215/Cdk5rap2, and Cep192 (*21*). This scaffold recruits γ-tubulin complexes for microtubule nucleation (*22–25*) and the motor protein Eg5/Kif11 to drive centrosome separation prior to nuclear envelope breakdown (NEBD) (*26, 27*). Following NEBD, a subset of newly formed microtubules composes anti-parallel bundles (interpolar microtubules) to slide the two spindle poles away, while another population of bundled microtubules (kinetochore fibers) captures the replicated sister chromatids to mediate their alignment and subsequent segregation (*28, 29*). The complex, multi-layered nature of the spindle assembly mechanism poses challenges for restoring its function via single-gene modification.

Recent studies have highlighted the E3 ubiquitin ligase TRIM37 as a master negative regulator of PCM assembly (*30–34*). TRIM37 mediates this process, at least in part, through promoting the degradation of Cep192 (*33, 34*). Remarkably, loss-of-function of TRIM37 facilitates the accumulation of functional PCM even in the absence of centrioles, enabling the efficient formation of a bipolar spindle in a cell lacking the centrosome. Consequently, TRIM37 deletion completely bypasses the requirement for the centrosome in proper mitotic control and stable proliferation across various human cell lines (*31, 33, 34*). This positions TRIM37-mediated PCM homeostasis as an attractive target for genetically engineering mitotic fidelity.

Here, we initially hypothesized that TRIM37 deletion could rescue mitotic fidelity in centrosome-depleted haploid cells. Paradoxically, we found that TRIM37 deletion was insufficient to rescue haploid stability due to a ploidy-specific structural fragility: mammalian haploid cells possess an intrinsically monopolar-prone spindle architecture. We demonstrate that this fragility stems from a haploidy-linked insufficiency in the absolute protein dosage of Cep192. By dissecting the molecular constraints of haploid spindle assembly, we define the Cep192–Aurora A–Eg5 axis as a critical bottleneck for haploid stability. Leveraging these insights in a genome-wide CRISPR-activation (CRISPRa) screen, we discovered versatile genetic targets that durably stabilize the human haploid state, illuminating the scaling principles of spindle assembly and establishing spindle polarity control as an actionable target for bioengineering.

## Results

### Limited effect of TRIM37 deletion on the stability of the haploid state in HAP1

Previous studies revealed that TRIM37 deletion eliminates the requirement of centrosomes in bipolar spindle formation in various human cell lines (*31, 33, 34*). Therefore, as a potential approach for engineering stable haploids, we tested whether TRIM37 deletion improves mitotic fidelity and chromosome stability in the haploid state. We used a human rare near-haploid cell line, HAP1, which is widely used as a model for haploid genetics and studies of haploidy-associated cellular defects (*1, 7, 13, 19, 35–38*). We first compared spindle organization between WT and TRIM37-KO haploid HAP1 cells by immunostaining microtubules, centrioles, and centrosomes (labeled by anti-α-tubulin, CP110, and PCNT, respectively; Fig. 1A; see also Fig. S1A for confirmation of TRIM37 deletion by immunoblotting). Consistent with our previous observations, 30% of WT haploids in mitosis manifested centrosome reduction (Fig. 1B) (*13, 20*). Most (97%) of these WT haploids with centrosome reduction possessed monopolar spindles (Fig. 1C). Centrosome reduction occurred in an equivalent proportion (30%) of TRIM37-KO haploids in mitosis (Fig. 1B), suggesting that TRIM37 deletion does not improve the centrosome retention in the haploid state.

**Fig. 1:**
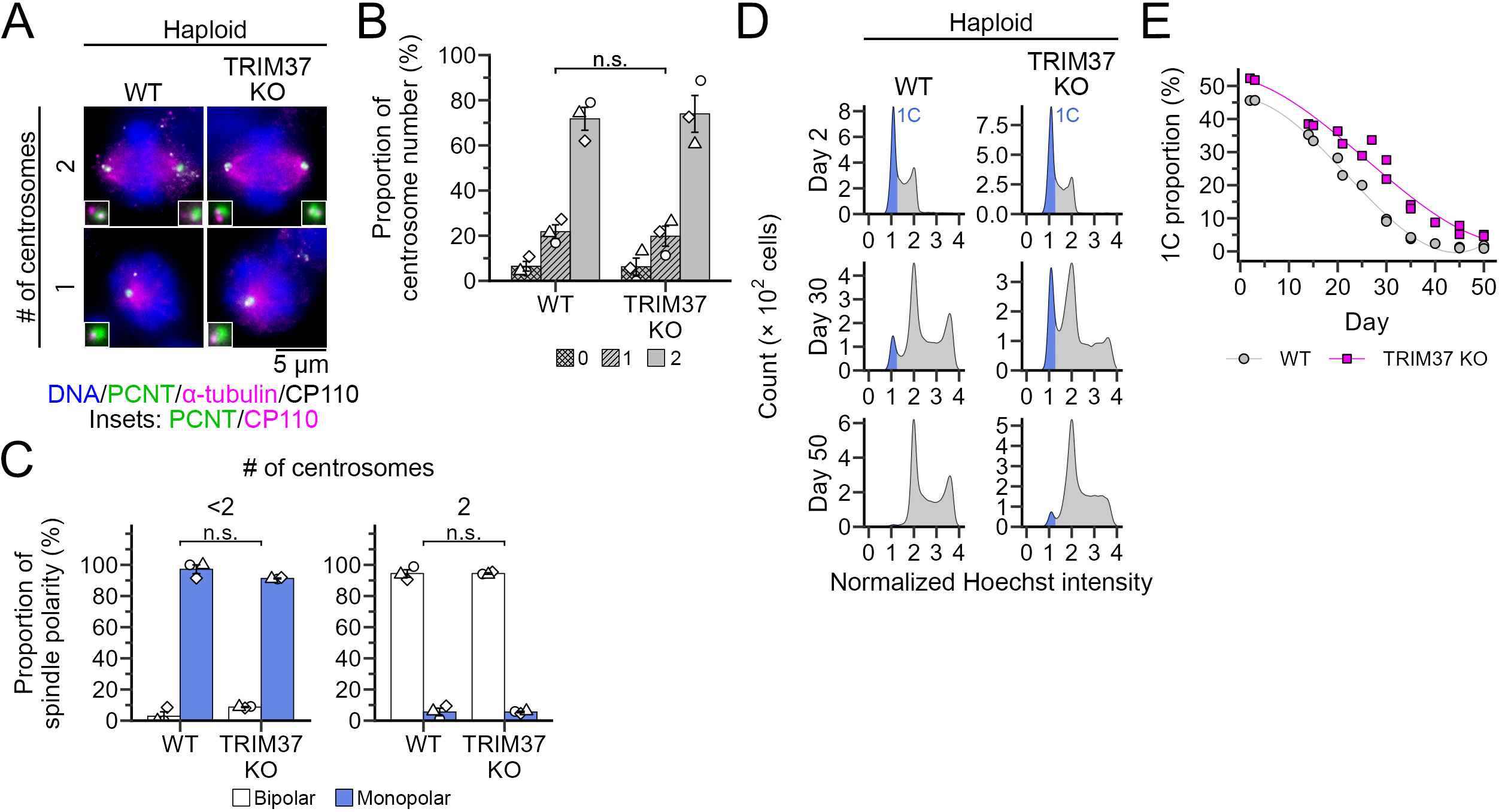
Limited effect of TRIM37 deletion on the stability of the haploid state in HAP1. **(A)** Immunostaining of α-tubulin, PCNT, and CP110 in WT or TRIM37-KO haploids in mitosis. DNA was stained with DAPI. Insets: 1.8× images of centrosomes. **(B, C)** Proportion of centrosome number (B) and spindle polarity (C) in A. Mean ± SE of 3 independent experiments. At least 323 cells were analyzed for each condition. In C, samples were sorted by centrosome number. There is no statistical significant difference in the proportion of monopolar spindles between conditions (n.s.: not significant, the Brunner–Munzel test). **(D)** Flow cytometric analysis of DNA content in WT or TRIM37-KO haploid cell culture during consecutive passages. DNA was stained with Hoechst 33342. The labels on the plot (1C) are relative DNA amounts (C-value) corresponding to haploid G1. **(E)** Time course of 1C (haploid G1) proportion in D. Data from 2 independent experiments are shown.

Though slightly less frequent than in the WT background, a substantial proportion of spindles (91%) remained monopolar under centrosome reduction even in the TRIM37-KO background (Fig. 1A and C). Therefore, contrary to our expectations, TRIM37 deletion had a very limited effect on bipolar spindle formation under innate centrosome loss in haploids.

We next compared chromosome stability between WT and TRIM37-KO haploids during long-term culture using flow cytometric DNA content analysis (Fig. 1D). Consistent with our previous observations (*13*), haploid WT cells gradually lost their haploid population, which was mostly replaced by the diploidized population by 40 d during consecutive passages (Fig. 1D and E). While TRIM37 deletion delayed the progression of haploid-to-diploid conversion, its effect was relatively mild: The haploid population was mostly lost by 50 d, even in the TRIM37-KO background (Fig. 1E). Since these findings potentially reflect fundamental issues associated to the haploid state, we decided to address the factors that limited the effect of TRIM37 depletion on spindle bipolarization and chromosome stability in haploids.

### Cep192 insufficiency underlies the attenuated acentrosomal spindle assembly in haploids

In previous studies using different human cell lines, loss-of-function of TRIM37 enhanced acentrosomal bipolar spindle assembly by facilitating the accumulation of PCM proteins, particularly Cep192, at the spindle poles (*31, 33, 34*). Therefore, we tested Cep192 protein expression level between WT and TRIM37-KO backgrounds using immunoblotting (Fig. S1A and B). To assess the potential effects of ploidy differences, we analyzed both haploid and diploid versions of WT or TRIM37-KO HAP1 cells. We previously found that haploids had approximately half the total protein content of diploids (*13*). In the immunoblotting, the relative Cep192 amount per total protein (estimated by housekeeping GAPDH amount) was equivalent between haploids and diploids, either in the WT or TRIM37-KO background (Fig. S1B). This result suggests that the Cep192 level per cell (i.e., an absolute dosage) was roughly halved in haploids compared to diploids.

We next compared Cep192 accumulation at the spindle poles in WT or TRIM37-KO haploids and diploids using immunostaining (Fig. 2A and B). To avoid bias from haploidy-linked centrosome loss, we analyzed only spindle poles with a centrosome. In WT haploids, the Cep192 level at the spindle poles was significantly less compared to the diploid control (Fig. 2A and B). Consistent with the previous reports (*33, 34*), TRIM37-KO significantly increased Cep192 at the spindle poles in diploids. On the other hand, in haploids, TRIM37-KO resulted in only a mild increase in Cep192 at the poles, without statistical significance (Fig. 2A and B). As a result, the Cep192 level at the poles was significantly lower in haploids than in diploids, even in the TRIM37-KO background (Fig. 2A and B).

**Fig. 2:**
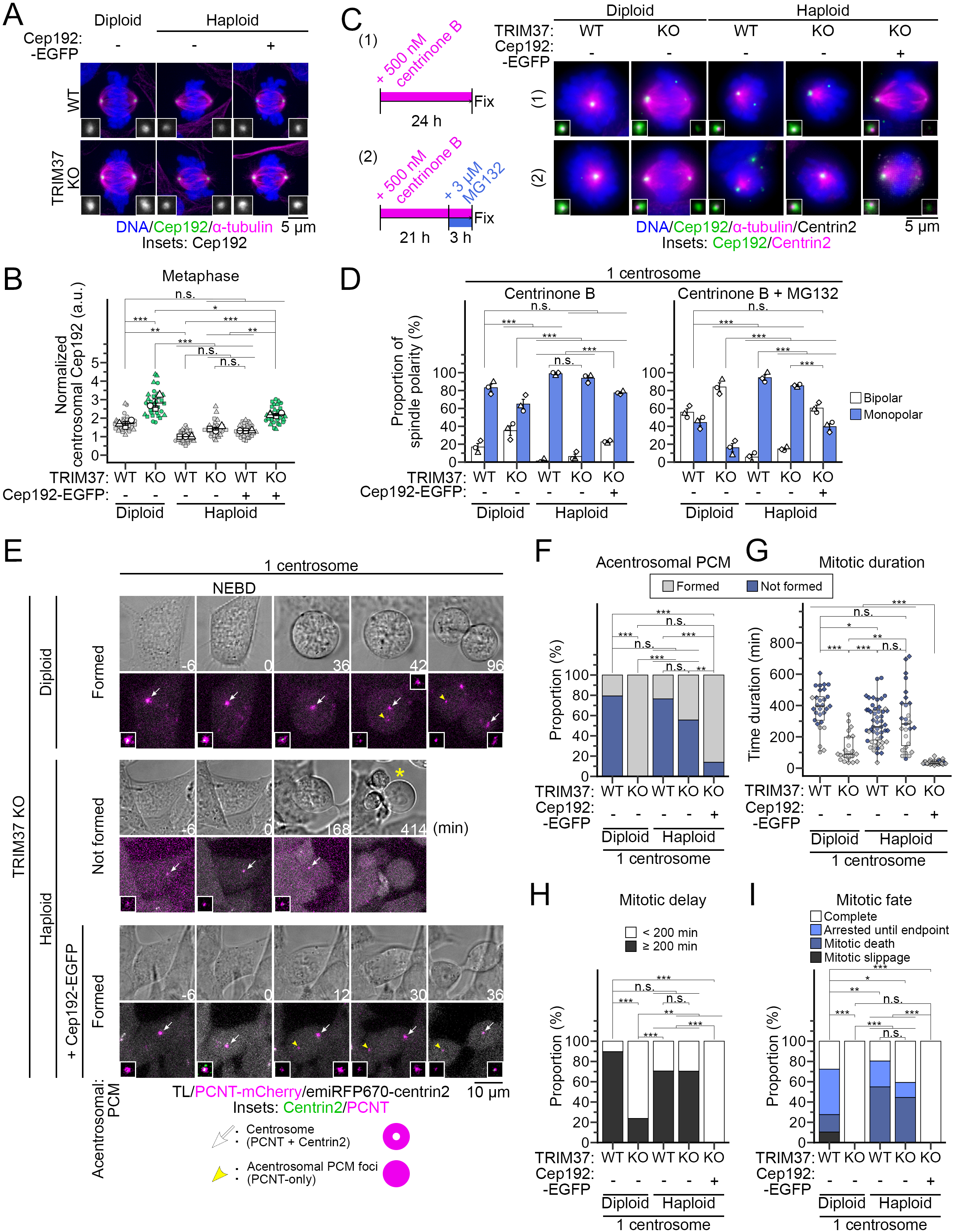
Cep192 insufficiency attenuates acentrosomal bipolar spindle assembly in haploids. **(A)** Immunostaining of α-tubulin and Cep192 in haploids or diploids with indicated backgrounds. DNA was stained with DAPI. Insets: 2.3× images of centrosomes. **(B)** Quantification of Cep192 immunostaining signals at the spindle poles in A (normalized to WT haploid). Mean ± SE of 3 independent experiments. All datapoints are also shown (at least 15 cells were analyzed for each condition). Asterisks indicate statistically significant differences between conditions (*p<0.05, **p<0.01, *** p<0.001, the Tukey–Kramer test). **(C)** Immunostaining of α-tubulin, Cep192, and centrin2 in haploids or diploids with indicated backgrounds, treated with the inhibitors as schematized on the left. DNA was stained with DAPI. Insets: 1.9× images of centrosomes. **(D)** Proportion of spindle polarity in C. Only cells with 1 centrosome (labeled by Cep192 with 1 or 2 centrin2 foci) were analyzed. Mean ± SE of 3 independent experiments. At least 91 cells were analyzed for each condition. Asterisks indicate statistically significant differences in the proportion of monopolar spindles between conditions (*** p<0.001, the Steel–Dwass test). **(E)** Time-lapse images of PCM reorganization during mitosis in haploids or diploids with indicated backgrounds, treated with centrinone B for 1 d. Live images were taken at 6-min intervals. PCM or centrioles were labeled with endogenous PCNT-mCherry or with transgenic emiRFP670-centrin2, respectively. White arrows or yellow arrowheads indicate centrosomes or acentrosomal PCM foci, respectively. The yellow asterisk indicates mitotic death. **(F)** Proportion of acentrosomal PCM foci formation in E. Asterisks indicate statistically significant differences between conditions (**p<0.01, *** p<0.001, the Fisher’s exact test with BH correction). **(G)** Mitotic duration (from NEBD to mitotic exit) in E. Asterisks indicate statistically significant differences between conditions (*p<0.05, **p<0.01, *** p<0.001, the DSCF test). **(H)** Frequency of mitotic delay in E. Asterisks indicate statistically significant differences between conditions (**p<0.01, *** p<0.001, the Fisher’s exact test with BH correction). **(I)** Proportion of mitotic fates in E. Asterisks indicate statistically significant differences between conditions (*p<0.05, **p<0.01, *** p<0.001, the Fisher’s exact test with BH correction). At least 21 cells pooled from 2 independent experiments were analyzed in F-I.

Given the ploidy-dependent difference in polar accumulation of Cep192, we tested whether the cellular effects of TRIM37 deletion differed between haploids and diploids. For this, we treated a Plk4 inhibitor, centrinone B, for 1 d, to induce acute centriole reduction and tested spindle organization in WT or TRIM37-KO haploids and diploids by immunostaining (Fig. 2C; note that centrosome loss does not naturally occur in diploid HAP1). To facilitate assessment of spindle bipolarization, we also compared spindle organization in the condition in which the metaphase-to-anaphase transition was blocked by MG132 treatment. At 1 d after administration of centrinone B, the centrosome number was reduced to 1 (i.e., 1 Cep192-positive centrosome with 1 or 2 centrin2-positive centrioles) in substantial proportions of haploids or diploids (see Fig. S2A for detailed centrosome number distribution). Both haploid and diploid WT cells with a single centrosome suffered a prominent spindle monopolarization (99% and 83%, respectively; Fig. 2C and D; see also Fig. S2B for spindle polarity in cells with different centrosome numbers). TRIM37-KO diploids frequently formed bipolar spindles even with a single centrosome (monopolar spindles were reduced to 65%; Fig. 2C and D), reproducing findings in other cell lines (*31, 33, 34*). In contrast, the majority of spindles in TRIM37-KO haploids (94%) remained monopolar when they had a single centrosome (Fig. 2C and D). The above trends were well conserved even under blockage of metaphase-anaphase transition: Whereas 84% TRIM37-KO diploids achieved bipolar spindle formation, 85% TRIM37-KO haploids remained monopolar (“+ MG132” in Fig. 2D). These data demonstrate that the limited effects of TRIM37 deletion on acentrosomal spindle assembly in haploid HAP1 cells are attributed to their haploidy, not the HAP1 cell background. Though we attempted to test spindle polarity in haploids lacking the centrosome after an extended centrinone B treatment, the drastic diploidization precluded us from observing haploid cells beyond 1 d (see below).

We reasoned that, if the reduced accumulation of Cep192 on the centrosomes were due to insufficient Cep192 dosage in haploids, artificial supplementation of it would improve its retention on the centrosomes. To address this idea, we introduced the Cep192-EGFP transgene into WT or TRIM37-KO haploids by stable cDNA integration (Fig. S1A and B). The transgene expression resulted in approximately 1.5- to 2.0-fold increase in total Cep192 amount in either WT or TRIM37-KO haploids (compared to non-transfected controls; Fig. S1B). Cep192-EGFP expression also increased the accumulation of Cep192 at the spindle poles, particularly significantly in the TRIM37-KO background (Fig. 2A and B). Moreover, supplementation of Cep192 drastically improved the retention of bipolar spindle in TRIM37-KO haploids with a single centrosome under centrinone B treatment (Fig. 2C, D, and S2B). These data indicate that haploidy-linked Cep192 insufficiency limits the effects of TRIM37 deletion on the enhancement of acentrosomal bipolar spindle assembly in the haploid state.

To gain more insights into the effects of Cep192 dosage on the dynamics of PCM and spindle pole organization under centriole reduction, we conducted live imaging of transgenic emiRFP670-centrin2 and mCherry-tagged endogenous PCNT (labeling centrioles and PCM, respectively) under centrinone B treatment (Fig. 2E). In a large proportion (> 70%) of WT haploids or diploids with a single centriole, PCM accumulated only to a single focal structure containing the centriole (Fig. 2E and F). These cell populations tended to suffer drastic mitotic arrest (> 200 min; Fig. 2G and H), often resulting in mitotic death, or mitotic slippage (mitotic exit without chromosome segregation; Fig. 2I). In a small proportion of these cells, the PCM focus split into two or another focus emerged, leading to bipolar cell division, as previously observed in HeLa cells (Fig. 2F) (*39*). TRIM37-KO in diploids, but not in haploids, strongly facilitated the emergence of PCM focus (Fig. 2E and F), restoring the ability of the cells to undergo bipolar division under centriole reduction (Fig. 2G-I). Cep192 supplementation in TRIM37-KO haploids drastically improved bipolar PCM accumulation by facilitating the emergence of PCM foci (Fig. 2E and F), thereby completely resolving mitotic delay and failure (Fig. 2G-I). These results demonstrate that insufficient Cep192 dosage is the primary factor limiting the potency of haploids in establishing bipolar PCM accumulation during acentrosomal spindle assembly.

### TRIM37-KO with Cep192 supplementation enables long-term haploid stabilization

We wished to understand how the ploidy-dependent difference in the potency of acentrosomal spindle assembly affected chromosome stability under centrosome reduction. For this, we tested the effects of 4-d consecutive centrinone B treatment on DNA contents in WT or TRIM37-KO haploids and diploids using flow cytometry (Fig. 3A). A large proportion of WT diploid or all WT haploids underwent doubling of their DNA contents, with substantial loss of their original G1 populations (2C or 1C, respectively) under centrinone B treatment (Fig. 3A and B). This result indicates that the unsolvable monopolar spindle defect facilitates whole-genome doubling under centrosome loss.

**Fig. 3:**
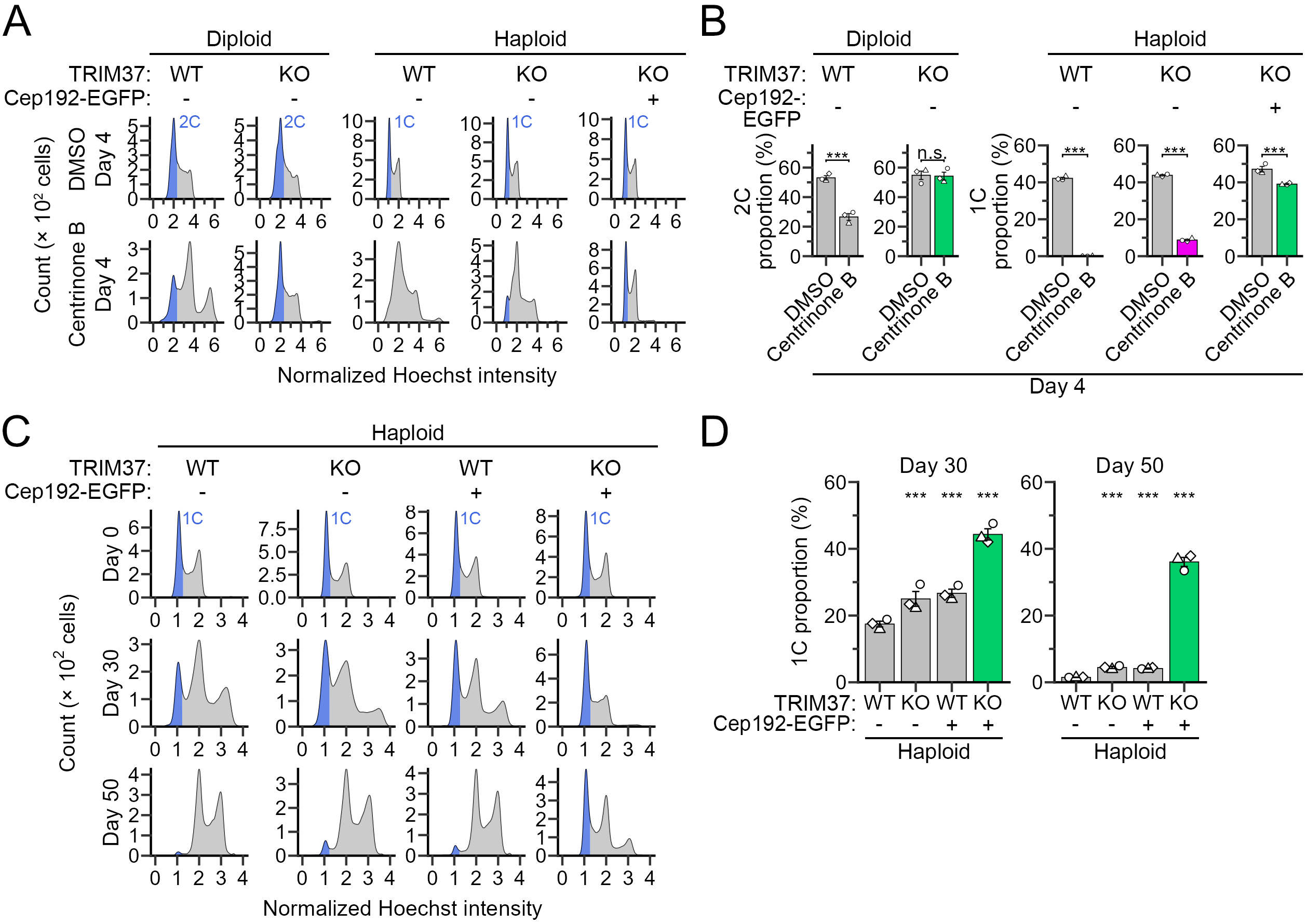
Cep192 supplementation drastically restores chromosome stability in TRIM37-KO haploids. **(A, C)** Flow cytometric analysis of DNA content in cells with indicated backgrounds, treated with or without centrinone B for 4 d (A), or during consecutive passages (C). DNA was stained with Hoechst 33342. **(B, D)** Frequency of 2C or 1C populations (for diploids or haploids, respectively) in A (B) or C (D) after the indicated duration of passages. Time course of 1C populations in C is shown in Fig. S2C. Mean ± SE of 3 independent experiments. Asterisks indicate statistically significant differences between conditions (B) or from haploid WT control (D) (***p<0.001, the Brunner–Munzel test (B) and the Steel test (D)).

TRIM37-KO completely resolved chromosome instability caused by centrinone B in diploids, with their DNA content kept equivalent to that in vehicle control (Fig. 3A and B). In contrast, TRIM37-KO improved chromosome stability only slightly in haploids under centrinone B treatment (Fig. 3A and B). Therefore, consistent with the haploidy-specific attenuation of acentrosomal spindle assembly (Fig. 2), the effect of TRIM37 deletion on chromosome stability was largely restricted in the haploid state. However, Cep192 supplementation in TRIM37-KO haploids nearly completely restored their chromosome stability under centrinone B treatment (Fig. 3A and B). This data further supports the idea that insufficient Cep192 dosage is the primary cause limiting the integrity of acentrosomal spindle assembly and the stability of the haploid state under centrosome loss.

We next asked whether restoring acentrosomal spindle assembly was sufficient to overcome naturally occurring haploid instability. To this end, we compared the dynamics of DNA content during long-term culture between WT and TRIM37-KO haploids, with or without Cep192 supplementation (Fig. 3C and D). Either TRIM37-KO or Cep192 supplementation singly had limited effects on haploid stability: While it modestly extended haploid G1 lifetime, most cells became diploids by 50 d. In contrast, the combination of TRIM37-KO and Cep192 supplementation resulted in a drastic improvement of the long-term stability of the haploid state (Fig. 3C and D): More than 72% of the original haploid G1 population was retained after 50 d, and haploid cells were still present in the culture even after 80 d (Fig. S2C). This finding demonstrates that the attenuation of spindle bipolarization mechanisms stemming from Cep192 insufficiency, in combination with haploidy-linked centrosome loss, is a fundamental cause of haploid instability in human somatic cells. The above data also indicate that genetic modification of spindle bipolarization mechanisms is an effective approach for stabilizing the naturally unstable haploid state in human cells.

### Cep192 insufficiency attenuates centrosome separation by the prophase pathway to retard spindle bipolarization in haploids

The above findings prompted us to address whether the haploidy-linked Cep192 insufficiency also affects mitotic control in the presence of a normal number (i.e., 2) of centrosomes. When cells possess 2 centrosomes, the initial step of bipolar spindle assembly is centrosome separation during prophase (referred to as the prophase pathway) (*40–42*). Therefore, we analyzed the dynamics of centrosome separation before and after NEBD using live imaging of mCherry-tagged endogenous PCNT and transgenic H2B-EGFP (to label centrosomes and chromosomes, respectively) in haploids and diploids (Fig. 4A, S3A, and B). All diploids (18 cells from 2 independent experiments) initiated centrosome separation during prophase and separated them over 4.0 μm by NEBD (Fig. 4B and S3B), demonstrating that the prophase pathway functions in the diploid state in HAP1. After NEBD, the inter-centrosomal distance continued to increase, reaching more than 8.8 μm within 15 min (Fig. S3B). In haploids, the progression of centrosome separation before NEBD was substantially retarded, with a substantial proportion (31%) hardly separating their centrosomes during prophase (with inter-centrosomal distance less than 4.0 μm at NEBD; Fig. 4A, B, and S3B). The centrosome separation kinetics were slower in haploids than in diploids, even when comparing only cell populations in which inter-centrosomal distance reached over 4.0 μm at NEBD (Fig. 4C). In haploids that failed to separate centrosomes over 4.0 μm by NEBD, the centrosomes remained unseparated (less than 1.7 μm in distance) for 15 min after NEBD (Fig. 4C and D). These data demonstrate that haploids are intrinsically inefficient in centrosome separation during prophase, leading to the innate retardation in spindle bipolarization.

**Fig. 4:**
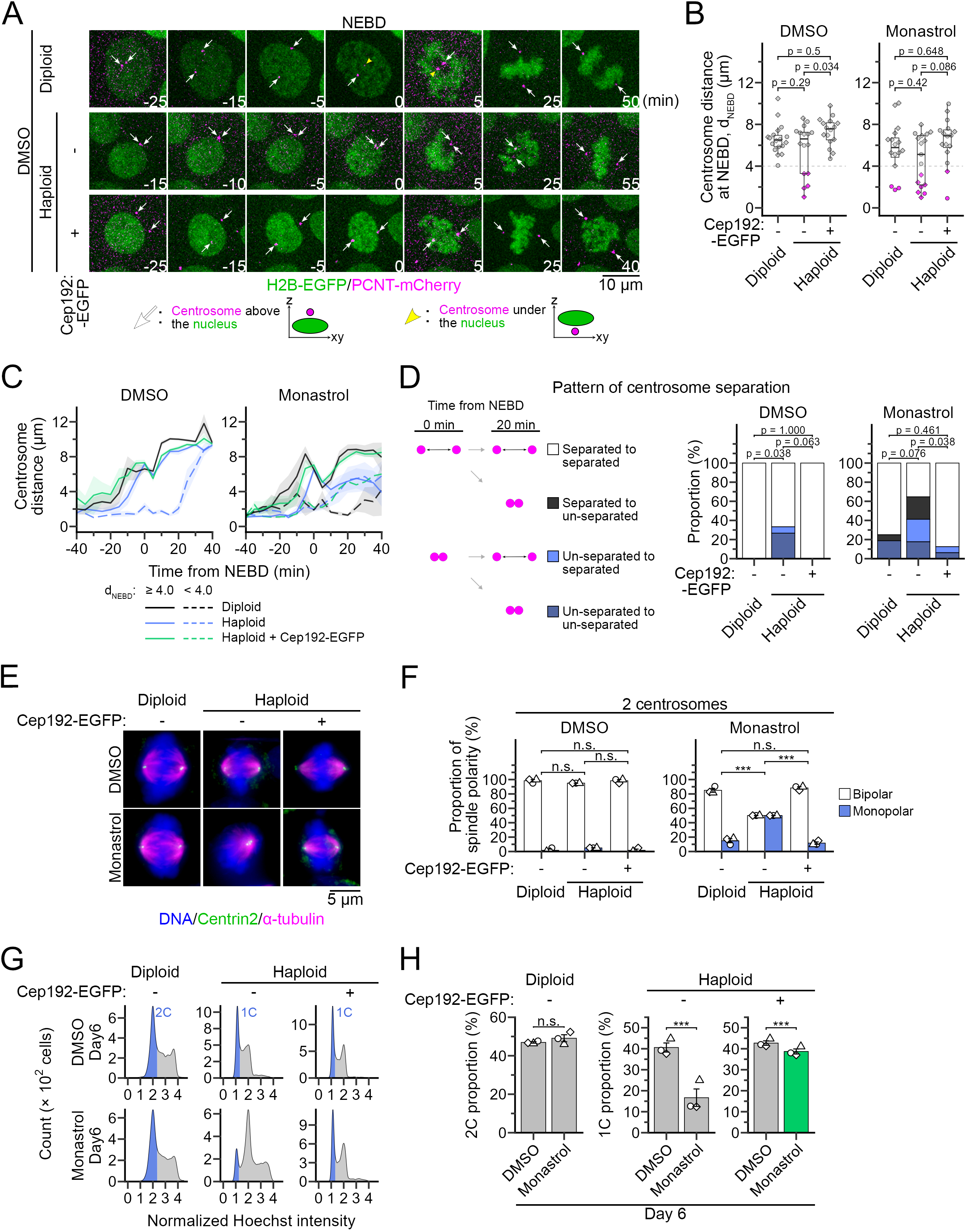
Haploidy-linked deficiency in spindle bipolarization in the presence of the centrosomes. **(A)** Time-lapse images of mitotic progression in haploids or diploids with indicated backgrounds. Centrosomes or chromosomes were labeled with endogenous PCNT-mCherry or with transgenic H2B-EGFP, respectively. Images of monastrol-treated cells are in Fig. S3A. Live images were taken at 5-min intervals. **(B)** Inter-centrosomal distance at NEBD (*d_NEBD_*) in A. At least 16 cells, pooled from 2 independent experiments, were analyzed for each condition (p-values from the DSCF test are shown). Box plots indicate the median, the 25^th^ and 75^th^ percentiles. **(C)** Time course of inter-centrosomal distance before and after NEBD in A. Cells with *d_NEBD_* of <4.0 μm and ≥4.0 μm were separately categorized. Mean ± SE of at least 3 cells from 2 independent experiments for each condition, except for Cep192-supplemented haploids with monastrol treatment (for this sample, the mean of 2 cells from 2 independent experiments is shown). The unsorted time course data are shown in Fig. S3B. **(D)** A scheme (left) or proportion (right) of centrosome separation patterns in A. At least 16 cells pooled from 2 independent experiments were analyzed for each condition (p values of the DSCF test are shown). **(E)** Immunostaining of α-tubulin and centrin2 in haploids or diploids with indicated backgrounds, treated with or without 12.5 μM monastrol. DNA was stained with DAPI. **(F)** Proportion of spindle polarity in E. Only cells with 2 centrosomes (each centrosome labeled by 2 centrin2 foci) were analyzed. Mean ± SE of 3 independent experiments. At least 62 cells were analyzed for each condition. Asterisks indicate statistically significant differences in the proportion of monopolar spindles between conditions (*** p<0.001, the Steel–Dwass test). **(G)** Flow cytometric analysis of DNA content in haploid or diploid cells with indicated backgrounds, treated with or without 25 μM monastrol for 6 d. DNA was stained with Hoechst 33342. **(H)** Frequency of 1C or 2C populations in G. Mean ± SE of 3 independent experiments. Asterisks indicate statistically significant differences between conditions (***p<0.001, the Brunner–Munzel test).

Cep192 supplementation in haploids facilitated the centrosome separation during prophase, restoring the kinetics of inter-centrosomal distance increase to the diploid level (Fig. 4A, C, and S3B). As a result, all Cep192-EGFP haploids achieved centrosome separation over 4.0 μm by NEBD and further separation during prometaphase, completely resolving the delay in spindle bipolarization in haploids (Fig. 4B and D). These data suggest that the innate inefficiency in centrosome separation and spindle bipolarization in haploids is primarily due to insufficient Cep192 dosage.

### Cep192 insufficiency underlies the haploidy-linked sensitization to Eg5 inhibition

In a previous comparative pharmacological study, we found that Eg5 inhibitors had significantly greater efficacy in suppressing the proliferation of haploid HAP1 cells than diploid cells, for unknown reasons (*43*). Based on the above results, we examined whether the defective bipolar spindle assembly in haploids underlies their vulnerability to Eg5 inhibition. We compared spindle organization under modest Eg5 inhibition by a relatively low concentration (12.5 μM) of monastrol between haploids and diploids using immunostaining (Fig. 4E). Monastrol had the most evident selectivity against haploid proliferation at this concentration (*43*). To avoid biases caused by differences in centrosome number, we analyzed only cells with two centrosomes (as indicated by four centrioles). In vehicle control, most haploids and diploids had bipolar spindles (Fig. 4F). A large proportion (85%) of diploids kept bipolar spindles under monastrol treatment (Fig. 4F), demonstrating that this condition was permissive for bipolar spindle assembly for diploids. In contrast, spindles were drastically monopolarized under the same condition in haploids (with the proportion of bipolar spindles reduced to 50%; Fig. 4F), demonstrating the profound proneness of haploids to monopolar spindle defects. Consistent with the immunostaining result, live imaging of centrosome separation also revealed a haploidy-linked aggravation of the centrosome separation defect under low-dose monastrol treatment (Fig. S3A and 4B-D). In particular, under modest Eg5 inhibition, we observed that bipolar spindles once formed and then reverted to monopolar more frequently in haploids than in diploids (Fig. 4D). This observation indicates that haploids have fragility not only in the establishment of centrosome separation but also in its maintenance for bipolar spindle assembly (see below). Increased expression of Cep192 in haploids restored bipolar spindle formation to the diploid level under modest Eg5 inhibition (Fig. 4E and F), with a drastic resolution of the centrosome separation defect (Fig. S3A and 4B-D). These results demonstrate that the haploidy-linked vulnerability to Eg5 inhibition is primarily attributed to Cep192 insufficiency.

We also compared the effects of the modest Eg5 inhibition on chromosome stability in haploids and diploids (Fig. 4G and H). Consistent with their haploidy-selective effects on mitotic spindles, modest Eg5 inhibition severely accelerated the diploidization of haploids but did not affect DNA content in diploid control (Fig. 4G and H). Importantly, supplementation of Cep192 almost entirely cancelled the Eg5 inhibitors-induced acceleration of diploidization in haploids. The above data collectively indicate that haploids’ vulnerability to Eg5 inhibition reflects their innate fragility in spindle bipolarization due to Cep192 insufficiency, potentially compromising chromosome stability independently of centrosome loss.

### Cep192 insufficiency limits centrosomal Eg5 recruitment for the prophase pathway in haploids

Since the primary defect caused by Cep192 insufficiency was an inefficient prophase pathway (Fig. 4), we further addressed how Cep192 dosage influenced centrosome separation in haploids. During prophase, Eg5 becomes concentrated at the centrosomes and plays an essential role in separating them (*26, 27, 41, 44*). Therefore, we compared Eg5 accumulation at the prophase centrosomes between haploids and diploids using immunostaining (Fig. 5A; cells with 2 centrosomes were analyzed). In haploids, the Eg5 accumulation level at the centrosomes was only 51% of that in diploids (Fig. 5A and B), suggesting the reduced functionality of Eg5 in centrosome separation.

**Fig. 5:**
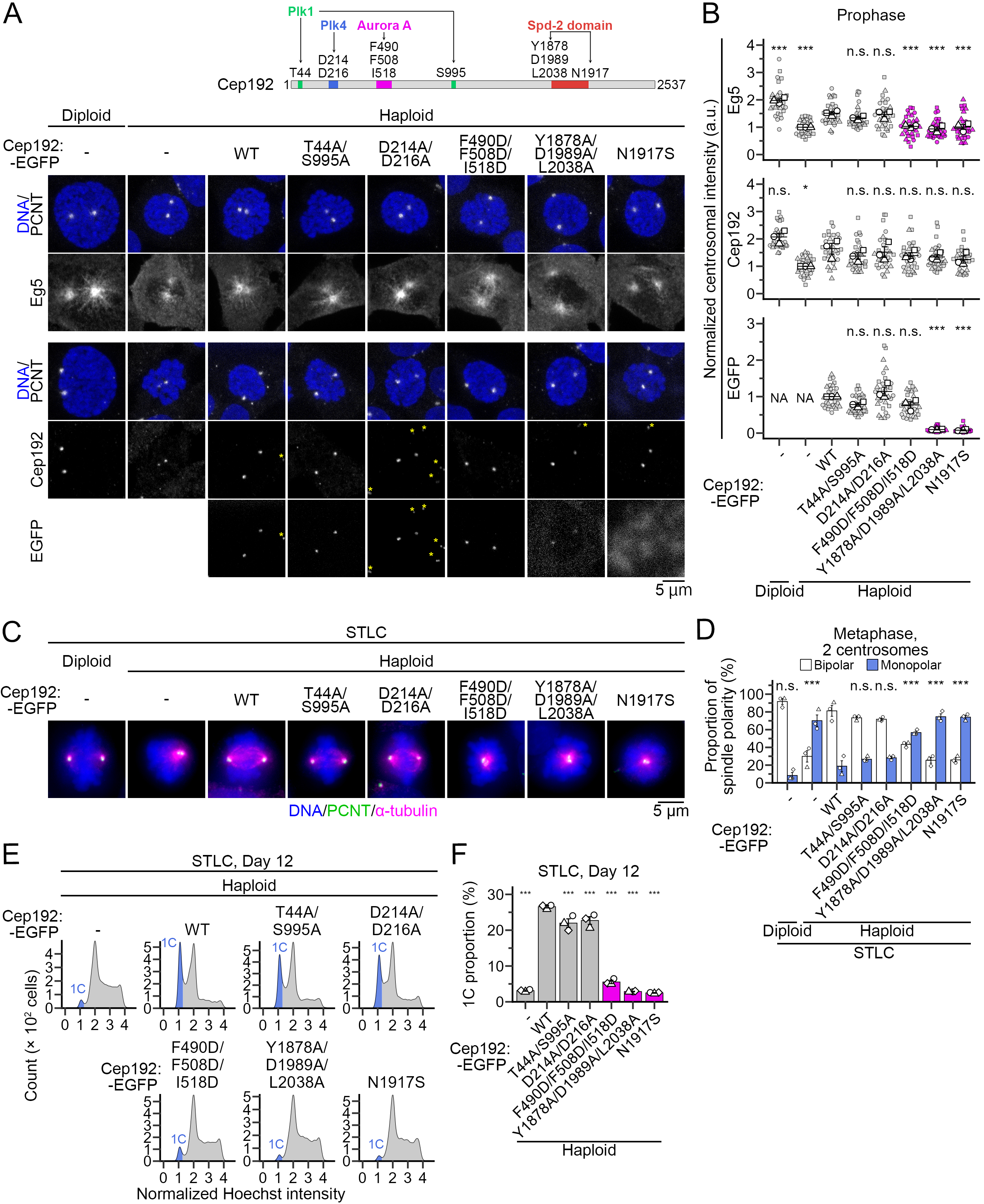
Cep192 supplementation restores centrosomal accumulation of Eg5 via interaction with Aurora A in haploids. **(A)** Immunostaining of Eg5 or Cep192, with PCNT in prophase haploids or diploids with indicated backgrounds. The scheme at the top depicts the mutations introduced into the Cep192-EGFP transgenes tested. DNA was stained with DAPI. Prophase cells were identified based on DAPI staining patterns. Asterisks indicate signals in neighboring cells. **(B)** Quantification of normalized Eg5 or Cep192 immunostaining signals or transgenic EGFP signals at the spindle poles in A. Mean ± SE of 3 independent experiments. All datapoints are also shown (at least 15 cells were analyzed for each condition). Asterisks indicate statistically significant differences from the haploid Cep192 (WT)-EGFP condition (*p<0.05, *** p<0.001, the Dunnett’s test). **(C)** Immunostaining of PCNT and α-tubulin in haploids or diploids with indicated backgrounds, treated with 300 nM STLC for 6 h. DNA was stained with DAPI. **(D)** Proportion of spindle polarity in C. Only cells with 2 centrosomes were analyzed. Mean ± SE of 3 independent experiments. At least 63 cells were analyzed for each condition. Asterisks indicate statistically significant differences in the proportion of monopolar spindles from the haploid Cep192 (WT)-EGFP condition (*** p<0.001, the Steel test). **(E)** Flow cytometric analysis of DNA content in haploids with indicated backgrounds, treated with 300 nM STLC for 12 d. DNA was stained with Hoechst 33342. **(F)** Frequency of 1C populations in E. Mean ± SE of 3 independent experiments. Asterisks indicate statistically significant differences from the haploid Cep192 (WT)-EGFP condition (*** p<0.001, the Steel test).

Cep192 supplementation by Cep192-EGFP expression substantially increased the centrosomal accumulation of Eg5 in haploids (to 76% of diploid level; Fig. 5A and B). Therefore, Cep192 insufficiency limits the centrosomal level of Eg5 in haploids, potentially explaining their inefficient centrosome separation during prophase.

Cep192 consists of multiple domains that mediate centrosomal accumulation of Cep192 itself, as well as the recruitment and activation of key mitotic kinases, Aurora A and Plk1, to organize PCM assembly (*24, 45–47*). To specify the aspects of Cep192 function relevant to the haploidy-linked defects, we established haploid lines expressing Cep192-EGFP that harbored mutations on various domains, in addition to endogenous Cep192 (Fig. 5A and B; see also Fig. S4A for immunoblotting). We introduced mutations on Aurora A- or Plk1-binding domain that abolished the interaction with the corresponding protein (F490D/F508D/I518D or T44A/S995A, respectively) (*48, 49*). We also introduced mutations on conserved amino acids in the Spd-2 domain (SP2D), equivalent mutations of which abolished centrosomal accumulation of Spd-2, a fly Cep192 orthologue, in fly embryos (Y1878A/D1989A/L2038A or N1917S) (*45*). We also tested mutations in the Plk4-binding domain (D214A/D216A) to interfere with Cep192-mediated recruitment of Plk4 for centriole duplication during interphase (*50*). All mutant transgenes were expressed at a level equivalent to that of Cep192 (WT)-EGFP (Fig. S4A and B).

We first tested the effects of transgene expression on centrosomal accumulation of Eg5 in prophase (Fig. 5A). Consistent with the report in fly embryos, Cep192 (Y1878A/D1989A/L2038A)-EGFP or Cep192 (N1917S)-EGFP failed to robustly accumulate on the centrosomes, diffusively localized in the cytoplasm (Fig. 5A). Therefore, the role of SP2D in mediating Cep192 accumulation to the centrosomes, found in flies (*45*), is conserved in human cells. Cep192 (Y1878A/D1989A/L2038A)-EGFP or Cep192 (N1917S)-EGFP expression failed to restore centrosomal Eg5 accumulation in haploids (Fig. 5B). Though all other mutants accumulated at the centrosomes as efficiently as Cep192 (WT)-EGFP (Fig. 5A and B), their effects on Eg5 recruitment were different: Whereas expression of Cep192 (T44A/S995A)-EGFP or Cep192 (D214A/D216A)-EGFP increased Eg5 at the centrosomes as efficiently as Cep192 (WT)-EGFP, Cep192 (F490D/F508D/I518D)-EGFP completely failed to restore Eg5 accumulation (Fig. 5B).

We also tested the effects of expression of these Cep192 mutants on spindle polarity and chromosome stability under modest Eg5 inhibition (treated with 300 nM S-trityl-L-cysteine (STLC); Fig. 5C-F, and S4C). Consistent with their effects on Eg5 recruitment, Cep192 (T44A/S995A)-EGFP and Cep192 (D214A/D216A)-EGFP improved spindle bipolarity (Fig. 5C and D) and retention of haploid DNA content (Fig. 5E and F), similar to Cep192 (WT)-EGFP. In contrast, Cep192 (F490D/F508D/I518D)-EGFP, Cep192 (Y1878A/D1989A/L2038A)-EGFP, and Cep192 (N1917S)-EGFP failed to improve either spindle bipolarity (Fig. 5C and D) or haploid stability (Fig. 5E and F), compared to the non-transfected control. These data collectively indicate that supplemented Cep192 restores Eg5 recruitment, bipolar spindle assembly, and the stability of the haploid state through its accumulation at centrosomes and interaction with Aurora A: These aspects of Cep192 function are likely key determinants of haploid stability.

### Loss of redundancy in the maintenance of spindle bipolarity in haploids

In various mammalian cell lines, Eg5 function becomes dispensable once bipolar spindles are assembled (*26, 51–53*). However, in the above live imaging, haploids often underwent bipolar-to-monopolar reversion under modest Eg5 inhibition (Fig. 4D), suggesting their fragility in maintaining spindle bipolarity. To further address this possibility, we arrested haploids or diploids at metaphase by treating them with MG132 for 90 min. After MG132 treatment, all haploids and diploids possessed bipolar spindles (Fig. 6A and B; only cells with two centrosomes were analyzed). These pre-arrested metaphase cells were then treated with MG132 and a high concentration (100 μM) of monastrol for 90 min (Fig. 6A). Consistent to the previous reports in other cell lines (*52, 53*), a large proportion (79%) of diploids maintained bipolar spindles after monastrol treatment (Fig. 6B). In contrast, spindles reverted to monopolar at a significantly higher frequency (54%) in haploids than in diploids after monastrol treatment (Fig. 6B; only cells with two centrosomes were analyzed). This result indicates that the mechanisms that maintain preformed bipolar spindles are attenuated in the haploid state.

**Fig. 6:**
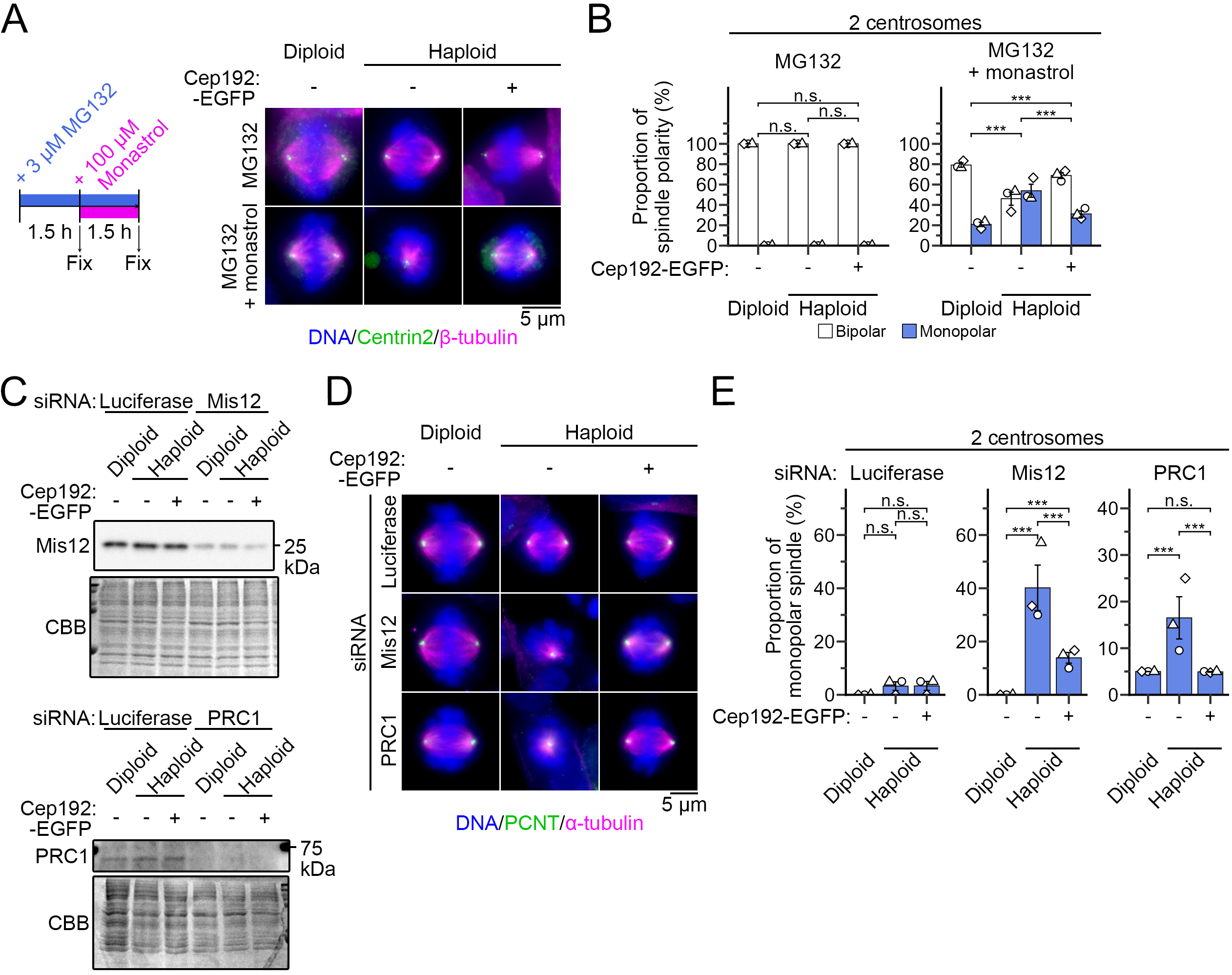
Loss of redundancy in maintenance of spindle bipolarity in haploids. **(A)** Immunostaining of β-tubulin and centrin2 in haploid or diploid cells with indicated backgrounds, treated with the inhibitors as schematized on the left. DNA was stained with DAPI. **(B)** Proportion of spindle polarity in A. Only cells with 2 centrosomes (each centrosome labeled by 2 centrin2 foci) were analyzed. Mean ± SE of 3 independent experiments. At least 65 cells were analyzed for each condition. Asterisks indicate statistically significant differences in the proportion of monopolar spindles between conditions (*** p<0.001, the Steel–Dwass test). **(C)** Immunoblotting of Mis12 (top) or PRC1 (bottom) in RNAi-treated haploid or diploid cells. CBB-stained total protein was quantified as the loading control. **(D)** Immunostaining of α-tubulin and PCNT in RNAi-treated haploid or diploid cells with indicated backgrounds. DNA was stained with DAPI. **(E)** Frequency of spindle polarity in D. Only cells with 2 centrosomes were analyzed. Mean ± SE of 3 independent experiments. At least 60 cells were analyzed for each condition. Asterisks indicate statistically significant differences between conditions (*** p<0.001, the Steel–Dwass test).

Bipolar spindles consist mainly of two distinct, but closely associated, microtubule populations: kinetochore fibers and interpolar microtubules, both of which contribute to the maintenance of their bipolarity (*28, 29, 40, 53–55*). Therefore, we next investigated the effects of selective perturbation of these microtubule populations on spindle morphology. We blocked the formation of kinetochore fibers or interpolar microtubules by depleting a kinetochore protein (hMis12) or an interpolar microtubule bundling protein (PRC1), respectively (Fig. 6C) (*56, 57*). In diploids, disorganization of either microtubule structure did not cause drastic spindle defects, suggesting that either of kinetochore- or interpolar microtubule-dependent mechanisms is dispensable for spindle bipolarization (Fig. 6D and E). In contrast, either depletion of Mis12 or PRC1 caused a significant increase in monopolar spindles in haploids (Fig. 6D and E; only cells with two centrosomes were analyzed). Therefore, in the haploid state, both microtubule populations are indispensable for spindle bipolarization, demonstrating the loss of their redundancy in maintaining spindle bipolarity. Interestingly, supplementing Cep192 in haploids substantially restored spindle bipolarity either under monastrol treatment after metaphase arrest (Fig. 6A and B) or the above RNAi treatments (Fig. 6D and E). These results suggest that Cep192 insufficiency also underlies the attenuation of spindle bipolarity maintenance in haploids.

### A genome-wide CRISPR-activation screen identifies gene targets for improving haploid stability

The above results demonstrate that haploid instability can be overcome by supplementation with a single gene, once appropriate gene targets are identified. This idea prompted us to conduct a genome-wide CRISPR-mediated transcriptional activation (CRISPRa) screen (*58*) to identify more genes whose activation could improve the stability of the haploid state. We wished to identify genes that stabilize the haploid state by improving the robustness of bipolar spindle assembly. Therefore, based on the observation that a low-dose Eg5 inhibitor accelerated diploidization of haploids (Fig. 4G), we designed the screen to identify genes that stabilize the haploid state under modest Eg5 inhibition (Fig. 7A; see Materials and methods for details). Haploid HAP1 cells stably expressing S.p. dCas9-VP64-p65-Rta (dCas9-VPR) (*59*) were infected with a lentiviral human CRISPRa guide RNA (gRNA) library (Calabrese P65-HSF) (*60*) and cultured for 6 d in the presence or absence of 300 nM STLC (Fig. 7A and S5A). Then, cell populations with DNA contents of 1C or 4C (corresponding to haploid G1 or diploid G2/M, respectively) were purified by flow cytometry after fixation and PI staining, and sequences of gRNAs retained in these populations were analyzed (Fig. 7A, S5B, and C; see Table S1 for all gRNA counts). As references, we also analyzed gRNA sequences derived from cell populations before drug administration (Pre-treatment) or from vehicle control (DMSO in Fig. 7A). To ensure the generality of the screen results, we also conducted the same CRISPRa screen in parallel using eHAP cells, a fully-haploidized derivative of HAP1 (treated with or without 400 nM STLC for 8 d) (*61*).

**Fig. 7:**
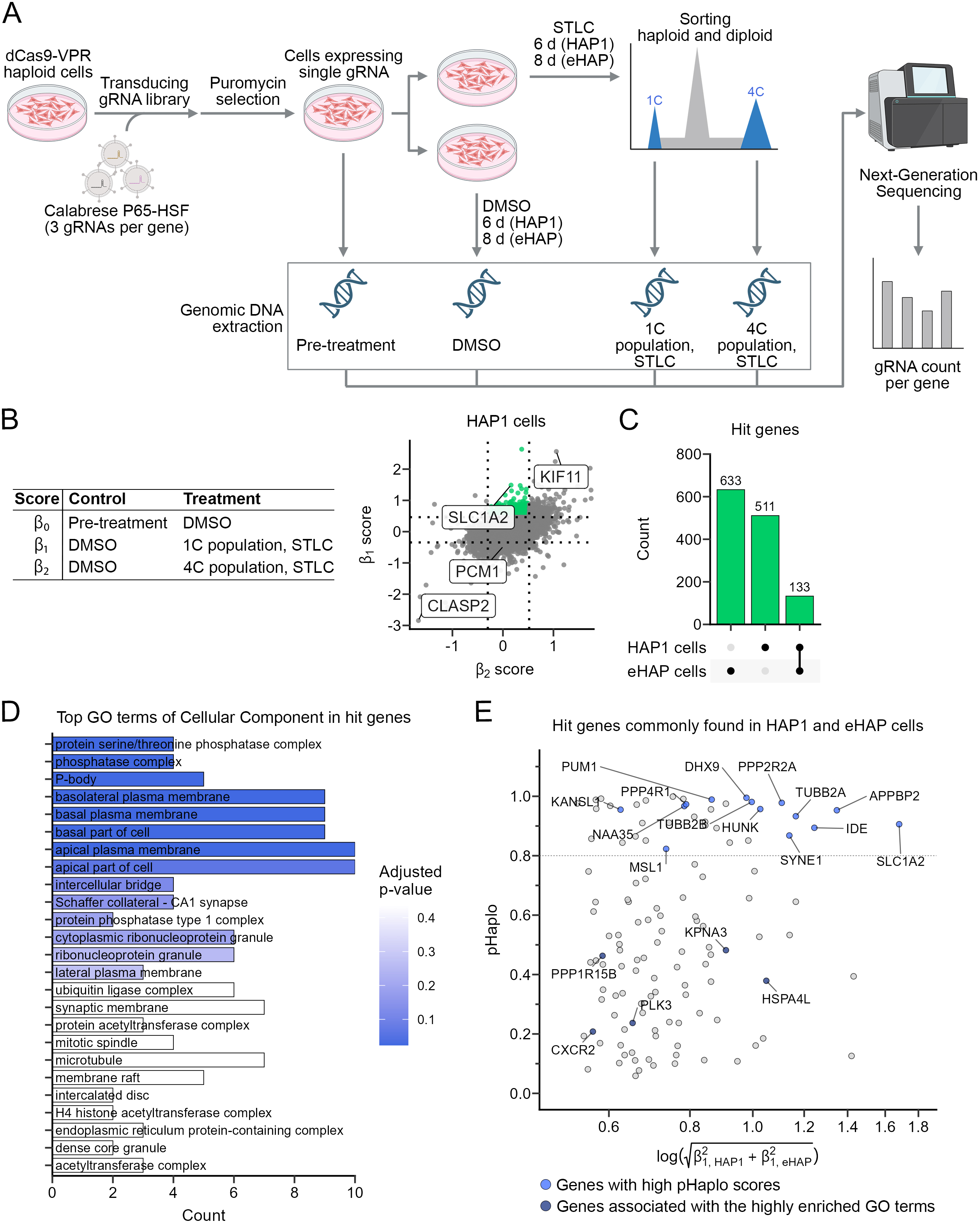
A genome-wide CRISPRa screen for haploid-stabilizing genes. **(A)** A scheme of CRISPRa screen to identify supplementation target genes for stabilizing the haploid state under modest Eg5 inhibition. **(B)** Comparison targets for each β score (left) and a 2-D scatter plot of β_1_ vs β_2_ scores in HAP1 (right). Hit genes selectively concentrated in the haploid G1 population under modest Eg5 inhibition were indicated in green. gRNA count profiles of highlighted genes are exemplified in Fig. S6A. **(C)** An upset plot of hit genes uniquely or commonly found in two haploid cell lines. **(D)** Gene ontology (GO) enrichment analysis for hit genes. **(E)** A 2-D plot of pHaplo score vs the logarithm of the dot product between β_1_ scores in HAP1 and eHAP cells. Data for 133 hit genes are shown. Candidate genes selected for further investigation were highlighted in blue or navy blue.

To specify the genes whose increase improved the stability of the haploid state, we identified gRNA targets significantly concentrated in haploid G1, but not in diploid G2/M, populations compared to vehicle control populations (633 or 511 genes in HAP1 or eHAP, respectively, with 133 genes commonly found in these cells; Fig. 7B, C, S6A-C, and Table S2). While Eg5 itself was found in the group of genes enriched in both 1C and 4C populations (*KIF11*; Fig. 7B and S6C), we decided to include it in the follow-up experiments as an important target of the Cep192-Aurora A signaling axis (*47*). A Gene Ontology (GO) enrichment analysis of genes selectively enriched in the 1C population (marked in green in Fig. 7B) revealed enrichment in pathways associated with protein phosphatases, membrane proteins, and so on (Fig. 7D). However, the computed connectivity among the candidates was overall sparse (Fig. S6D). Therefore, we decided to conduct follow-up analyses using a strategy distinct from the one focusing on specific gene pathways.

We reasoned that, among the candidates, genes with high dosage sensitivities would exhibit more pronounced biological effects when CRISPRa increased their expression levels. Based on this idea, we attempted to narrow down the candidates by selecting genes with higher expected dosage sensitivities. We graded the candidate genes based on their pHaplo scores, a metric that indexes the probability of haploinsufficiency, computed from an analysis of rare copy-number variants across patient genomes (Fig. 7E) (*62*). For the follow-up experiments, we selected 14 genes showing prominent enrichment in the 1C population with high pHaplo scores (highlighted in blue in Fig. 7E) and established haploid HAP1 cell lines stably expressing gRNAs targeting each candidate with dCas9-VPR. We also selected 5 genes associated with highly enriched GO terms, despite lower pHaplo scores (highlighted in navy blue in Fig. 7E).

We addressed whether CRISPRa-mediated upregulation of each candidate improved the retention of the haploid population under low-dose (300 nM) STLC treatment (Fig. 8A). Among the candidates, the upregulation of *KIF11*, *TUBB2A*, *SLC1A2*, *HUNK*, *IDE*, or *SYNE1* significantly improved the retention of the haploid G1 population after 6-d culture in the presence of STLC, while that of other genes did not (Fig. 8A, B; see also Fig. S7A for confirmation of gene upregulation of these positive hits). Therefore, we selected these 6 genes as potential haploid-stabilizing factors and further tested the effect of their upregulation on spindle assembly in the presence of 300 nM STLC (Fig. 8C, D, and S7B; cells with 2 centrosomes were analyzed). Consistent with the previous result (Fig. 4E), non-transfected control haploids frequently formed monopolar spindles even in the presence of 2 centrosomes (Fig. 8C and D). However, upregulation of each of these 6 genes resulted in a significant improvement in spindle bipolarity in the presence of STLC (Fig. 8D). Therefore, we identified multiple upregulation targets that strengthen the spindle bipolarization mechanism in the haploid state, thereby improving haploid stability, at least under Eg5 inhibition.

**Fig. 8:**
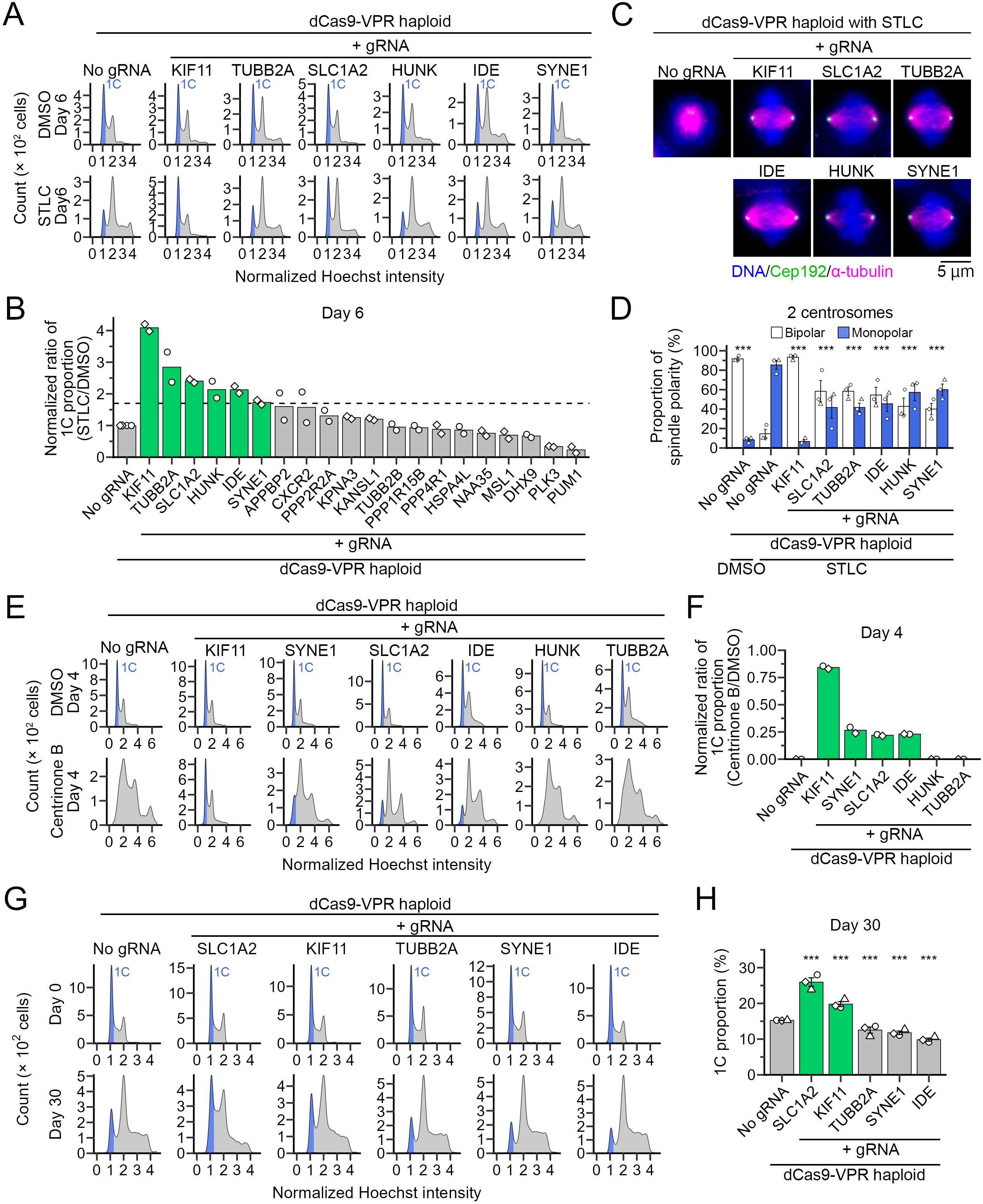
Supplementation of several genes resolves haploidy-linked spindle defects or chromosome instability. **(A)** Flow cytometric analysis of DNA content in haploid dCas9-VPR cells expressing gRNA for the indicated genes, treated with or without 300 nM STLC for 6 d. DNA was stained with Hoechst 33342. **(B)** Ratios of 1C populations after STLC treatment, relative to those in vehicle control for each cellular background, in A. Ratios were normalized to that of the gRNA-untransfected control. Mean of 2 independent experiments. Genes whose transcription activation increased the normalized 1C proportion above 1.7-fold of the untransfected control (indicated by the broken line) were selected for further investigation. **(C)** Immunostaining of α-tubulin and Cep192 in haploid dCas9-VPR cells expressing gRNA for the indicated genes, treated with 300 nM STLC. DNA was stained with DAPI. **(D)** Proportion of spindle polarity in C. Only cells with 2 centrosomes were analyzed. Mean ± SE of 3 independent experiments. At least 68 cells were analyzed for each condition. Asterisks indicate statistically significant differences in the proportion of monopolar spindles from the untransfected control treated with STLC (*** p<0.001, the Steel test). **(E, G)** Flow cytometric analysis of DNA content in haploid dCas9-VPR cells expressing gRNA for indicated genes, treated with centrinone B for 4 d (E), or during consecutive passages (G). DNA was stained with Hoechst 33342. **(F, H)** Frequency of 1C populations in E (F, normalized to that in vehicle control) or G (H). Mean of 2 independent experiments (F) or mean ± SE of 3 independent experiments (H). Asterisks indicate statistically significant differences from the untransfected control (*** p<0.001, the Steel test).

We next addressed whether upregulation of the above genes also improves haploid stability under different perturbations. To test this, we assessed the stability of the haploid state under 500 nM centrinone B treatment (Fig. 8E and F). In the non-transfected control, originally haploid cells almost fully converted to diploids after 4-d culture in the presence of centrinone B (Fig. 8E and F). On the other hand, upregulation of *KIF11*, *SYNE1*, *SLC1A2*, or *IDE* significantly improved the retention of the haploid G1 population in the same condition (Fig. 8E and F), demonstrating their prominent effect on haploid stability even under centrosome reduction. This finding prompted us to examine whether upregulation of these genes can help extend the lifespan of the haploid state in long-term passages (Fig. 8G), where naturally occurring centrosome loss drives progressive diploidization (Fig. 1D and E) (*20*). Consistent with their effects on the spindle fidelity, upregulation of *SLC1A2* or *KIF11* resulted in a significant improvement in haploid retention after 30 d of consecutive culture (Fig. 8G and H). To test the sustainability of their effects on haploid stability, we size-sorted the remaining 1C populations from control, *SLC1A2*-, or *KIF11*-upregulated cell cultures and subjected them to an additional 37-d culture (Fig. S7C). Their haploid-stabilizing effects were preserved throughout another round of consecutive culture (Fig. S7C and D), demonstrating the long-lasting effects of gene supplementation on maintaining the haploid population. These results directly support the concept that enhancing the bipolar spindle assembly mechanism is an effective approach for engineering haploid stability and identify effective gene targets for this purpose.

## Discussion

Somatic haploid cells are promising resources for state-of-the-art bioengineering, but their persistent instability limits their versatility in vertebrate models. In this study, we demonstrate that human haploids suffer an innate, structural fragility in establishing and maintaining mitotic spindle bipolarity. These spindle defects are primarily attributed to Cep192 insufficiency in the haploid state (Figs. 2, 4, and 6). Supplementing Cep192 proves highly effective in improving mitotic fidelity and chromosome stability of the haploid state (Figs. 2, 3, and 5), orthogonally to previously characterized centrosome loss (*13*). By defining the mechanical bottlenecks of haploidy-linked chromosome instability, we further identified a suite of genetic targets for stabilization of haploidy in human cells.

Previous studies have revealed profound scalability of the spindle structure in response to cellular dimensional changes associated with development or ploidy increase (*63–65*). In contrast, our data suggest that, in the haploid state, the spindle structure falls into a “scaling paradox,” which cannot be resolved by its inherent flexibility. Since a haploid cell has about half the cytoplasmic volume of a diploid cell (*9, 13*), the overall concentration of Cep192 is estimated to be roughly constant across ploidies (Fig. S1A). However, our data indicate that a Cep192 concentration equivalent to that in diploids alone is insufficient to secure a functional level of Cep192 accumulation at centrosomes (Fig. 2). This may be because critical absolute amounts of PCM components are required to achieve their cooperative scaffolding or phase transition (*66–69*), at a level sufficient to support spindle fidelity. We propose that, in haploids, halved gene dosage reduces the absolute cytoplasmic Cep192 pool below this structural threshold. The lack of spindle rescue by Cep192 mutants, which are unable to mediate PCM scaffolding (Fig. 5) (*45*), also supports the above idea. A recent study in C. elegans embryos demonstrated a similar phenomenon, where the Cep192 orthologue Spd-2 acts as a limiting factor that scales centrosome size to cell volume during embryogenesis (*69*). Our findings extend this biophysical principle to human somatic cells, proving that the absolute quantity of Cep192 rigidly dictates spindle functionality and genome stability.

Through structure-function mapping, we isolated the Aurora A-binding domain of Cep192 as the indispensable node for haploid spindle rescue. While Cep192 regulates both Aurora A and Plk1 (*46, 47, 49*), mutations that abolish Aurora A interaction completely abrogated the rescue of Eg5 recruitment and establishment of spindle bipolarity (Fig. 5). This finding aligns with our CRISPRa screen results, where direct transcriptional activation of *KIF11* (Eg5) was sufficient to restore spindle integrity and stabilize haploidy (Fig. 8). Therefore, Cep192 insufficiency drives haploid instability primarily by restricting the local Aurora A activation required to load Eg5 onto mitotic centrosomes. Meanwhile, since Cep192 supplementation restored the maintenance of preformed bipolar spindles even under stringent Eg5 inhibition (Fig. 6), the increased Cep192 would also exert its effects through pathways other than Eg5, possibly through its role in facilitating microtubule generation (*22*), for spindle maintenance. Comparative ultrastructural analyses across different ploidy levels would provide more insights into the limited scalability of spindle structure in the haploid state.

The drastic improvement in mitotic fidelity through single-gene supplementation prompted us to perform a genome-wide CRISPRa screen to identify orthogonal haploid stabilizers. We identified multiple novel targets, whose involvement in mitotic control has not been reported to our knowledge, except for *TUBB2A* and *SYNE1* (*70, 71*). To better understand the restriction of mitotic control in the haploid state, elucidating the mechanisms of mitotic restoration through supplementation of these genes, especially the high-affinity glutamate transporter *SLC1A2* (EAAT2), is an important future research goal. While *SLC1A2* is typically studied in the context of neuroprotection and excitotoxicity (*72, 73*), its profound effect on stabilizing the haploid spindle in non-neural HAP1 cells suggests its new roles in mitotic fidelity control. We speculate that *SLC1A2* upregulation may improve spindle fidelity through facilitating tubulin polyglutamylation (*74, 75*). Interestingly, the introduction of a negative charge on the C-terminal tail of tubulin through this modification is proposed to increase microtubule association and modulate the processivity of key mitotic factors, including Eg5 and PRC1 (*76, 77*). Therefore, *SLC1A2* may increase the intracellular glutamate pool required to hyper-glutamylate microtubules, fortifying the spindle against collapse. Another possibility is that enhanced glutamate import by *SLC1A2* upregulation facilitates the TCA cycle as a primary anaplerotic substrate via conversion to α-ketoglutarate (75), thereby buffering the immense ATP demand for mitotic control. Dissecting how SLC1A2-mediated metabolic fluxes influence spindle biomechanics represents a fascinating avenue for future investigation.

The genes identified through the CRISPRa screen showed different degrees of haploid-stabilizing effects (Fig. 8). A clear limitation of gene supplementation via CRISPRa is the lack of quantitative control over gene upregulation. Therefore, a key task for future research is to determine the extent to which the stabilization effects of each gene can be improved through more elaborate dosage titration. Furthermore, since we identified many orthogonal targets, it will be intriguing to address whether synergistic effects arise from their combined upregulation.

Somatic haploidy causes pleiotropic defects in organismal functions (*9*). Therefore, the biological effects of the genome-wide gene dosage insufficiency would extend beyond mitosis to diverse cellular processes. The gene supplementation screen in a haploid background, such as those performed in this study, would provide unique opportunities to identify new dosage-dependent factors playing crucial roles in various cellular functions. Genetic engineering of mitotic control presented here successfully stabilized innately unstable haploid cells to a degree that allows them to be cultured and handled as effortlessly as standard diploid lines, thereby dramatically expanding the horizon for haploid cell technology in animal systems.

## Materials and Methods

### Cell culture and cell line establishment

HAP1 or eHAP cells (RRID: CVCL_Y019 or CVCL_5J33, respectively; from Haplogen GmbH) were cultured in Iscove’s Modified Dulbecco’s Medium (IMDM; Wako Pure Chemical Industries) supplemented with 10% fetal bovine serum (FBS) and 1× antibiotic-antimycotic solution (AA; Sigma-Aldrich) at 37°C with 5% CO_2_. Culture dishes were pre-coated with 10% collagen Type I-C (Wako, 637-00773) diluted in phosphate-buffered saline (PBS). Transgenic cell lines were established by transfecting cells with plasmids encoding the corresponding transgenes using Nucleofector 2b (Lonza), followed by selection of positive clones with the appropriate antibiotics. The TRIM37-KO HAP1 line (RRID: CVCL_TU16) was purchased from Horizon. Cell lines used in this study are listed in Table S3. All experiments were performed with mycoplasma-free cells (tested using the MycoStrip mycoplasma detection kit; InvivoGen). Haploid HAP1 or eHAP lines were maintained by size-based cell sorting, and diploid HAP1 lines were established as previously described (*13, 43*).

### Plasmids, compounds, and antibodies

The plasmid vectors constructed or used in this study, and the primers used for their construction are listed in Table S4 and Table S5, respectively. Plasmids for endogenous mCherry tagging to PCNT (*39*) were gifts from Takumi Chinen and Daiju Kitagawa (University of Tokyo). Compounds and antibodies used in this study are listed in Table S6. The siRNAs used in this study are 5’-GGACAUUUUGAUAACCUUUtt-3’ (Mis12; DNA in lowercase), 5’-AAAUAUGGGAGCUAAUUGGGA-3’ (PRC1), and 5’-CGUACGCGGAAUACUUCGAtt-3’ (luciferase). siRNA transfection was performed using Lipofectamine RNAiMAX (Thermo Fisher Scientific).

### Flow cytometry

All flow cytometric analyses, except for those in the CRISPRa screen, were conducted using a JSAN desktop cell sorter (Bay Bioscience). For live-cell DNA content analyses, cells were trypsinized using 0.05% trypsin-EDTA solution (Wako) and stained with 10 μg/ml Hoechst 33342 (Dojindo) for 15 min at 37°C. For fixed-cell DNA content analyses, cells were fixed with cold 70% ethanol overnight at 4°C, stained with 1 μg/ml Hoechst 33342 in the presence of 5 μg/ml ribonuclease (Nippon Gene) for 1 h at 25°C. The stained cells were filtered through a cell strainer and applied to the cell sorter.

The Hoechst or PI intensities from individual cells were plotted into a histogram and subjected to Kernel density estimation to identify 1C and 2C peaks. The DNA content of individual cells was then normalized so that the 2C peak was set to 2. To define the ranges for 1C and 2C populations, we set the standard deviation (SD) for each population as follows. The SD for the 2C population (*SD_2C_*) was set to 0.15 in normalized units. Then, the SD of 1C populations (*SD_1C_*) was defined using the equation below:

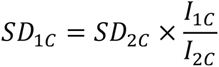

where *I_1C_* or *I_2C_* was normalized DNA content at 1C or 2C peak, respectively. The range of the individual population was defined as the 1st to 99th percentile of a normal distribution centered on the peak, with the corresponding SD, then used to calculate the 1C or 2C proportion. The R script used for the automated cell-proportion analysis above is provided in Supplemental Material S1.

### Metaphase synchronization

To assess cells’ ability to maintain preformed bipolar spindle under perturbations, asynchronous cell cultures were treated with 3 μM MG132 for 90 min at 37°C with 5% CO_2_. “MG132” control cultures were fixed and immunostained at this step to assess the efficiency of metaphase arrest and spindle polarity after MG132 treatment. Remaining cultures were subjected to a second treatment with 3 μM MG132 and 100 μM monastrol for another 90 min, then fixed for immunostaining.

### Immunofluorescence staining

For immunostaining of centrin2 and CP110, cells were fixed with 100% methanol at -20°C for 10 min, and treated with BSA blocking buffer (150 mM NaCl, 10 mM Tris-HCl, pH 7.5, 5% BSA, and 0.1% Tween 20) at 25°C for 30 min. For immunostaining of α-tubulin, β-tubulin, Cep192, Eg5, and PCNT, cells were fixed with 3.2% paraformaldehyde in PBS for 10 min, permeabilized with 0.5% Triton-X100 in PBS supplemented with 0.1 M glycine GPBS for 15 min at 25°C, and treated with blocking buffers (1% BSA in PBS for α-tubulin, β-tubulin, Cep192, and PCNT, and 3% donkey serum in PBS for Eg5). Fixed cell samples were then incubated with the first antibodies overnight at 4°C, followed by incubation with the fluorescence-conjugated secondaries overnight at 4°C. DNA was stained with 1 μg/mL DAPI. Following each treatment, cells were washed 3 times with PBS.

Fixed cells were observed under one of the following microscopes: A Ti2 microscope (Nikon) with a ×100 1.45 NA Plan-Apochromatic and Zyla4.2 sCMOS camera (Andor); A Ti microscope (Nikon) with a ×100 1.40 NA Plan-Apochromatic, and Nikon A1Rsi (Nikon).

### Live-imaging experiments

For live imaging, we seeded cells on an 8-well cover glass-bottom chamber (Iwaki or Zell-kontakt) coated with 10% collagen Type I-C (Wako, 637-00773) diluted in PBS, and replaced culture media with phenol red-free IMDM (Gibco) supplemented with 10% FBS and AA 2 h before starting imaging. For centrinone B or monastrol treatment, we administered either compound at 22 h or 2 h, respectively, before starting imaging.

Live cell imaging was conducted under either of the following microscopes at 37°C with 5% CO2: A TE2000 microscope with ×60 1.4 NA Plan-Apochromatic, CSU-X1, and iXon3; A Ti2-E microscope (Nikon) with a ×60 1.4 NA Plan-Apochromatic, and Nikon A1Rsi (Nikon).

### Quantitative analyses of imaging data

For spindle polarity analyses, spindle poles were visualized by immunostaining either the centrioles (centrin2 or CP110) or centrosomes (Cep192 or PCNT), except for Fig. 1A and 2C, where centrin2 and Cep192 were co-immunostained. In polarity analyses under centrinone B treatment, spindle poles with or without the centrosome were distinguished based on the presence or absence of centrin2-positive signals. Spindles with only one pole or two poles, whose distance was less than 4 μm, were defined as monopolar. Interpolar distances were manually measured using the line tool in Image J (NIH).

For quantification of centrosomal accumulation of mitotic factors, immunostaining signals in centrosomal regions were measured from circular ROIs with diameters of 2.5 or 1.7 µm (for Eg5 or Cep192, respectively), manually drawn using the oval selection tool in ImageJ. For background correction, intensities of non-cell areas were also measured manually and subtracted from the ROI intensities.

To analyze the dynamics of centrosome or PCM separation in live imaging, their three-dimensional positions were manually recorded at each time point using the Cell Counter tool in ImageJ.

### Immunoblotting

Cells were lysed in SDS-PAGE sample buffer (1.125% SDS, 35 mM Tris-HCl, pH 6.8, 11.25% glycerol, 5% 2-mercaptoethanol) or RIPA buffer (50 mM Tris–HCl, pH 7.6, 150 mM NaCl, 10 mM NaF, 10 mM b-glycerophosphate, 1% NP-40, 0.5% sodium deoxycholate and 0.1% SDS, protein inhibitor cocktail (cOmplete, Roche)) on ice, and then boiled for 5 min. Proteins separated by SDS-PAGE were transferred onto an Immun-Blot PVDF membrane (Bio-Rad). The membranes were blocked with 0.3% skim milk in Tween Tris-buffered saline (TTBS; 50 mM Tris, 138 mM NaCl, 2.7 mM KCl, and 0.1% Tween 20), incubated with the first antibodies overnight at 4°C, and incubated with horseradish peroxidase (HRP)-conjugated secondary antibodies for 1-3 h at 25°C. Each step was followed by 3 washes with TTBS. Signals were detected using the ezWestLumi plus ECL Substrate (ATTO, Tokyo, Japan) with a LuminoGraph II chemiluminescent imaging system (ATTO). Quantification of immunoblotting signals was conducted using the Gels tool in ImageJ.

### RT-qPCR

Approximately 0.5-1.0 × 10^6^ cells in asynchronous cultures were collected by trypsinization, and their total RNA was extracted using Nucleospin RNA kit (Macherey-Nagel). Reverse transcription was performed with 200 ng of total RNA in a 10 μL reaction using ReverTra Ace qPCR RT Master Mix (Toyobo). qPCR was then performed with 50 ng cDNA template and 500 nM forward and reverse primers using GeneAce SYBR^TM^ qPCR Mix II (Nippon Gene) on a Real-time PCR analysis system (Bio-Rad, CFX96). Each step was conducted according to the manufacturer’s instructions. We analyzed the data using the ΔΔCq method, with the mean Cq values of the β-actin gene used to calculate ΔCq. The primers used for qPCR are listed in Table S5.

### CRISPR activation screen

#### - Library preparation

Human Calabrese CRISPR Activation Pooled Library Set A (59) was a gift from David Root and John Doench (Addgene #92379). The library was packaged in Lenti-X293T cells by co-transfecting with pMDLg/pRRE, pRSV-Rev, and pMD2.G (gifts from Didier Trono; Addgene plasmids #12251, #12253, and #12259, respectively) (*78*), using X-tremeGENE 9 (6365809001, Roche). At 48 h after transfection, supernatants were collected. Then, fresh Dulbecco’s Modified Eagle’s Medium (Sigma-

Aldrich) containing 10% FBS (supplemented DMEM) was supplied to the cultures for another 24-h incubation, followed by supernatant collection. The collected supernatant was filtered through a 0.45 μm syringe filter and subjected to viral concentration using the Lenti-X Concentrator (Takara Bio) according to the manufacturer’s instructions. Precipitated viruses were then resuspended in supplemented DMEM in a volume equivalent to one-tenth of the original supernatant and frozen in aliquots.

Lentivirus titration was conducted using the Lentivirus Titer p24 ELISA Kit (GenScript) according to the manufacturer’s instructions. For determining the optimal viral titer, dCas9-VPR HAP1 or eHAP cells were transduced with serial dilutions of the lentiviral library using 8 μg/mL polybrene (Sigma-Aldrich) and subjected to 1 μg/mL puromycin selection. At 4 d after transduction, when all cells had died in the untransduced puromycin-treated culture, we counted living cells in all cultures using a LUNA-II Automated Cell Counter (Logos Biosystems) with 0.4% Trypan Blue (Sigma-Aldrich). Infection efficiencies at different viral dilutions were calculated as the rate of surviving cells relative to that in a non-puromycin-treated culture at the same dilutions. We selected the viral titer that gave approximately 25% infection efficiency (corresponding to a multiplicity of infection (MOI) of 0.2) for the following CRISPRa screen.

#### - CRISPRa screen for haploid stabilizing genes

dCas9-VPR HAP1 or eHAP cells were transduced with the lentiviral library at an MOI of ∼0.2 in the presence of 8 μg/mL polyberene. We transduced 60 × 10^6^ cells to achieve a representation of 5 cells per gRNA. From 2 d after transduction, cells were subjected to 1 μg/mL puromycin selection for 2 d or 3 d in HAP1 or eHAP cells, respectively. After the puromycin selection, 200 × 10^6^ cells were fixed with 70% ethanol at 4°C for gRNA sequencing (“Pre-treatment”). Remaining cells were subjected to further passages in the presence or absence of 300 nM STLC for 6 d or 400 nM STLC for 8 d (for HAP1 or eHAP, respectively) and fixed with 70% ethanol at 4°C. The fixed vehicle control cells were directly subjected to gRNA sequencing (“DMSO”). The fixed STLC-treated cells were further subjected to DNA staining using FxCycle PI/RNase staining solution (Invitrogen), followed by DNA content-based cell sorting using a Cell Sorter SH800S (SONY) for isolating haploid G1 and diploid G2/M populations (“1C population” and “4C population,” respectively).

Throughout the passages, cell numbers in each condition were kept at least 20 × 10^6^ cells, ensuring coverage of at least 200 cells per gRNA.

#### Sequencing

The genomic DNA was extracted from each fixed cell population using the Wizard Genomic DNA Purification Kit (Promega) according to the manufacturer’s instructions. The extracted genomic DNA was then subjected to gRNA amplification using NEBNext Ultra II Q5 (New England Biolabs) with primers listed in Table S5, followed by next-generation sequencing on the NovaSeq X Plus PE150 platform.

#### Data processing and β scores

The gRNA read count files for each cell type and experimental condition were generated from the raw CRISPR fastq files using the “count” function in the MAGeCK software (v0.5.9.5) (*79*). β scores (computed degrees of gene selection between comparative conditions) (*79*) for each gene were estimated from these read count files using the MAGeCK-MLE algorithm (*79*). Combinations of conditions for each β score are shown in Fig. 7B. The quality of the fastq files and count data was evaluated using FastQC (v0.12.1, Babraham Bioinformatics) and MAGeCK software, respectively (Fig. S5B and C).

#### Identification of genes accumulated in the haploid population

To exclude genes whose gRNA read counts fluctuated drastically during cell culture, regardless of the Eg5 inhibitory state, we filtered out genes whose *β*_0_ scores (gene selection in vehicle control relative to pre-treatment) exceeded the top and bottom 0.75% thresholds in both HAP1 and eHAP cells (Fig. S6B). From the remaining genes, we selected those whose *β*_1_ scores, but not *β*_2_ scores (gene selection in the STLC-treated 1C or 4C population, respectively, relative to the vehicle control) exceeded the top 5% threshold for candidate haploid-stabilizing factors.

#### Functional enrichment analysis

Gene ontology (GO) enrichment analysis of Cellular Component for the 133 identified genes from the CRISPR activation screen was conducted using the enrichGO function from the clusterProfiler (v4.14.6) (*80*). The network of identified genes and shared GO terms was visualized using the cnetplot function from the clusterProfiler.

### Statistical analysis

Analyses for significant differences between the two groups were conducted using the Brunner-Munzel test. Multiple group analyses were conducted using the Tukey-Kramer test, the Fisher’s exact test with the Benjamini-Hochberg multiple testing correction, the Steel-Dwass test, or the Dwass-Steel Critchlow-Fligner (DSCF) test. Multiple group analyses with common control were conducted using the Dunnett’s test and the Steel test. All statistical analyses were conducted in R software. Statistical significance was set at *p* < 0.05. *P*-values are indicated in figures or the corresponding figure legends.

## Acknowledgment

We are grateful to Takumi Chinen and Daiju Kitagawa for plasmids, Mithilesh Mishra for the arrangement of the international collaboration, Kiminori Nakamura and Tokiyoshi Ayabe for lending us their research equipment, Global Facility Center at Hokkaido University for the flow cytometer and real-time PCR system, and the Nikon Imaging Center at Hokkaido University for microscopes.

## Author Contributions

Conceptualization, K.Y, and R.U.; Methodology, K.Y., H.R.S, P.K., J.Z., and R.U.; Investigation, K.Y.; Formal Analysis, K.Y.; Resources, K.Y., H.R.S, P.K., J.Z., and R.U.; Supervision, J.Z., and R.U.; Writing – Original Draft, R.U.; Writing – Review & Editing, K.Y., J.Z., and R.U.; Funding Acquisition, K.Y., J.Z., and R.U.

## Funding and additional information

This work was supported by the Sasakawa Scientific Research Grant and Hokkaido University-Hitachi Joint Cooperative Support Program for Education and Research to Koya Yoshizawa, JSPS KAKENHI (Grant Numbers JP21K19244, JP22H04926, JP24K21956, JP24KK0139, and JP24K02017), the Princess Takamatsu Cancer Research Fund, the Orange Foundation, the Smoking Research Foundation, Daiichi Sankyo Foundation of Life Science, the Akiyama Life Science Foundation, the Hoansha Foundation, Sumitomo Electric Group CSR Foundation, and the Terumo Life Science Foundation to Ryota Uehara.

## Data and material availability

The data supporting this study’s findings are available from the corresponding author upon reasonable request.

---

**Fig. S1:**
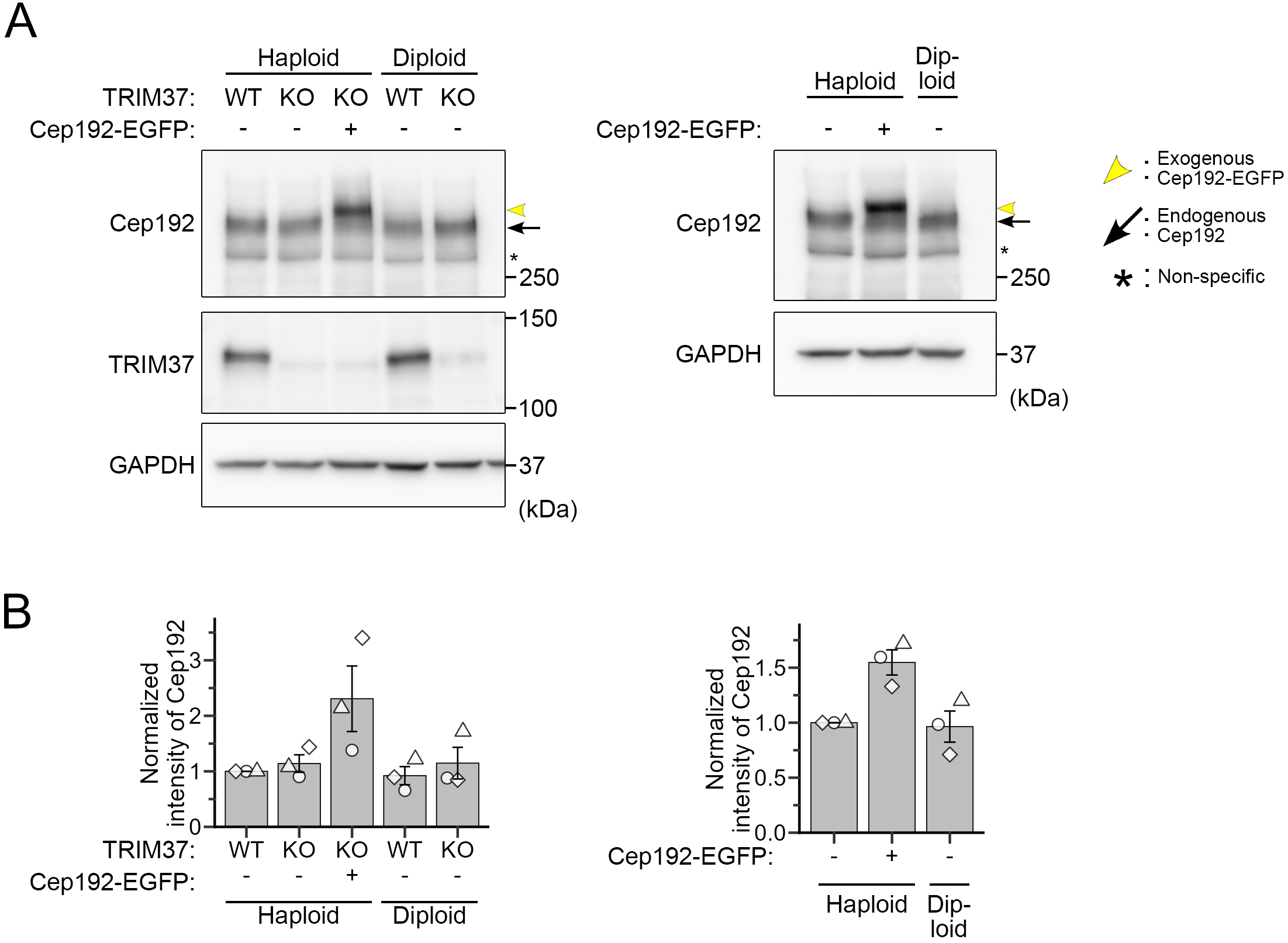
Immunoblotting analysis of TRIM37 or Cep192 expression in HAP1 cells. **(A)** Immunoblotting of TRIM37 or Cep192 in haploid or diploid cells with the indicated backgrounds. GAPDH was detected as a loading control. **(B)** Quantification of the relative intensity of Cep192 in A. Protein loading differences were corrected based on GAPDH signals.

**Fig. S2:**
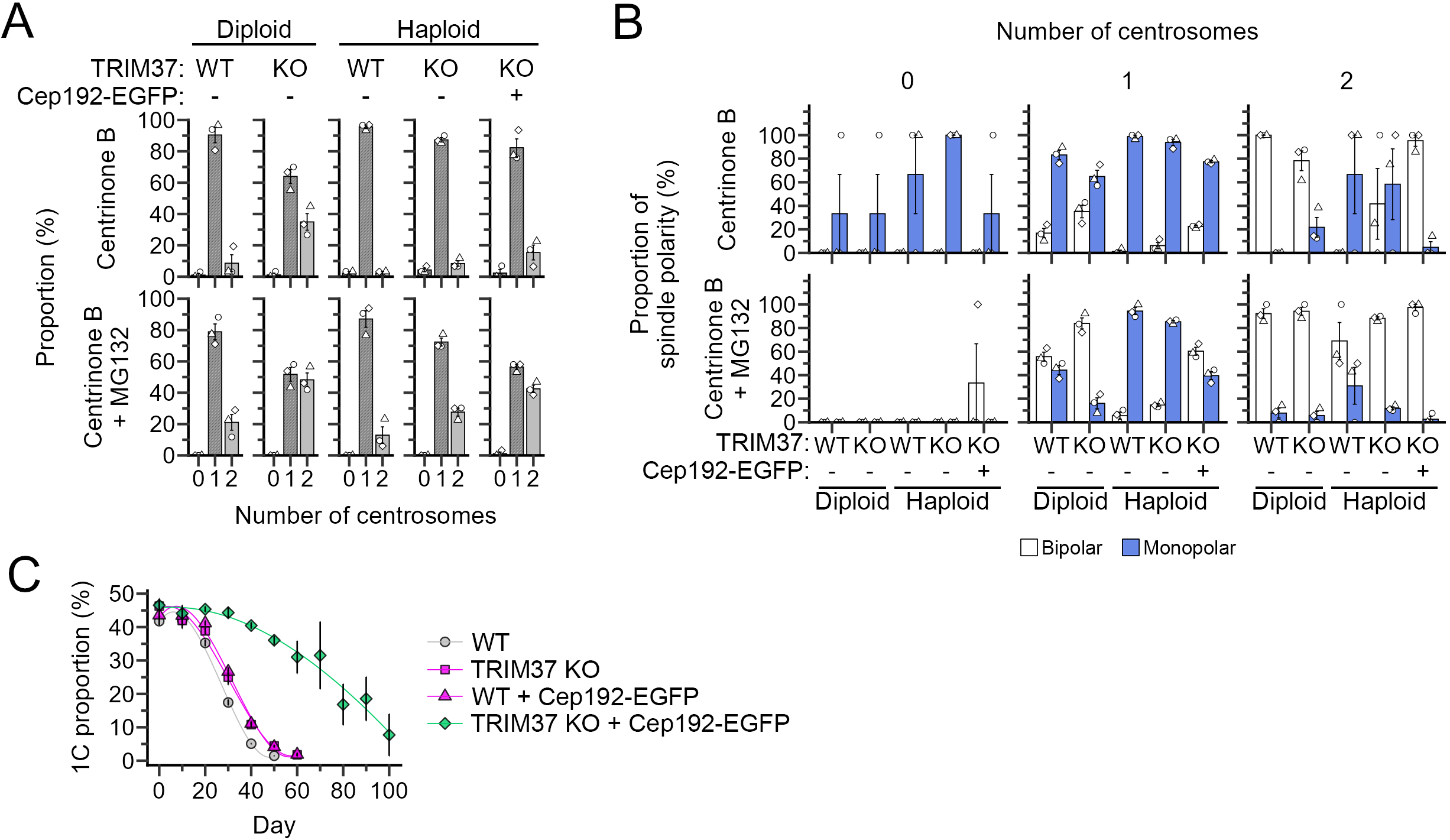
Effects of Cep192 supplementation on centrosome number, spindle polarity, and long-term stability in haploids. **(A)** Proportion of centrosome numbers in Fig. 2C and D. Mean ± SE of 3 independent experiments. At least 91 cells from 3 independent experiments were analyzed for each condition. **(B)** Proportion of spindle polarity in Fig. 2C, sorted by centrosome number. Mean ± SE of 3 independent experiments. At least 91 cells were analyzed for each condition. Identical data for the “1 centrosome” sample are shown in Fig. 2D. **(C)** Time course of 1C proportion in Fig. 3C. Mean ± SE of 3 independent experiments.

**Fig. S3:**
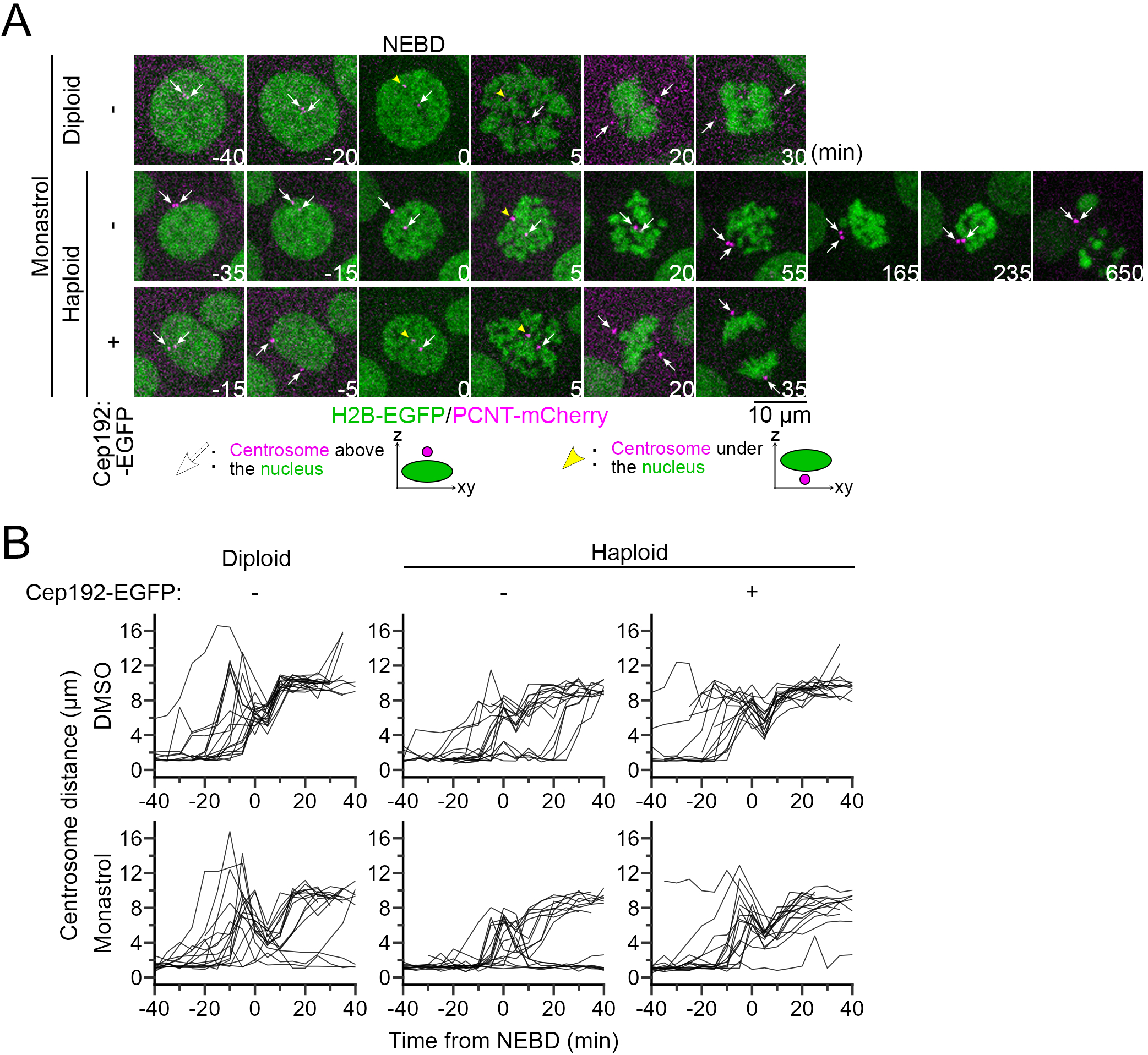
Live imaging of centrosome separation during mitosis. **(A)** Time-lapse images of mitotic progression in haploids or diploids with indicated backgrounds, treated with 12.5 μM monastrol. Centrosomes or chromosomes were labeled with endogenous PCNT-mCherry or with transgenic H2B-EGFP, respectively. **(B)** Time course of inter-centrosomal distance before and after NEBD in all individual cells in Fig. 4A or S3A. Data from at least 16 cells from 2 independent experiments are shown.

**Fig. S4:**
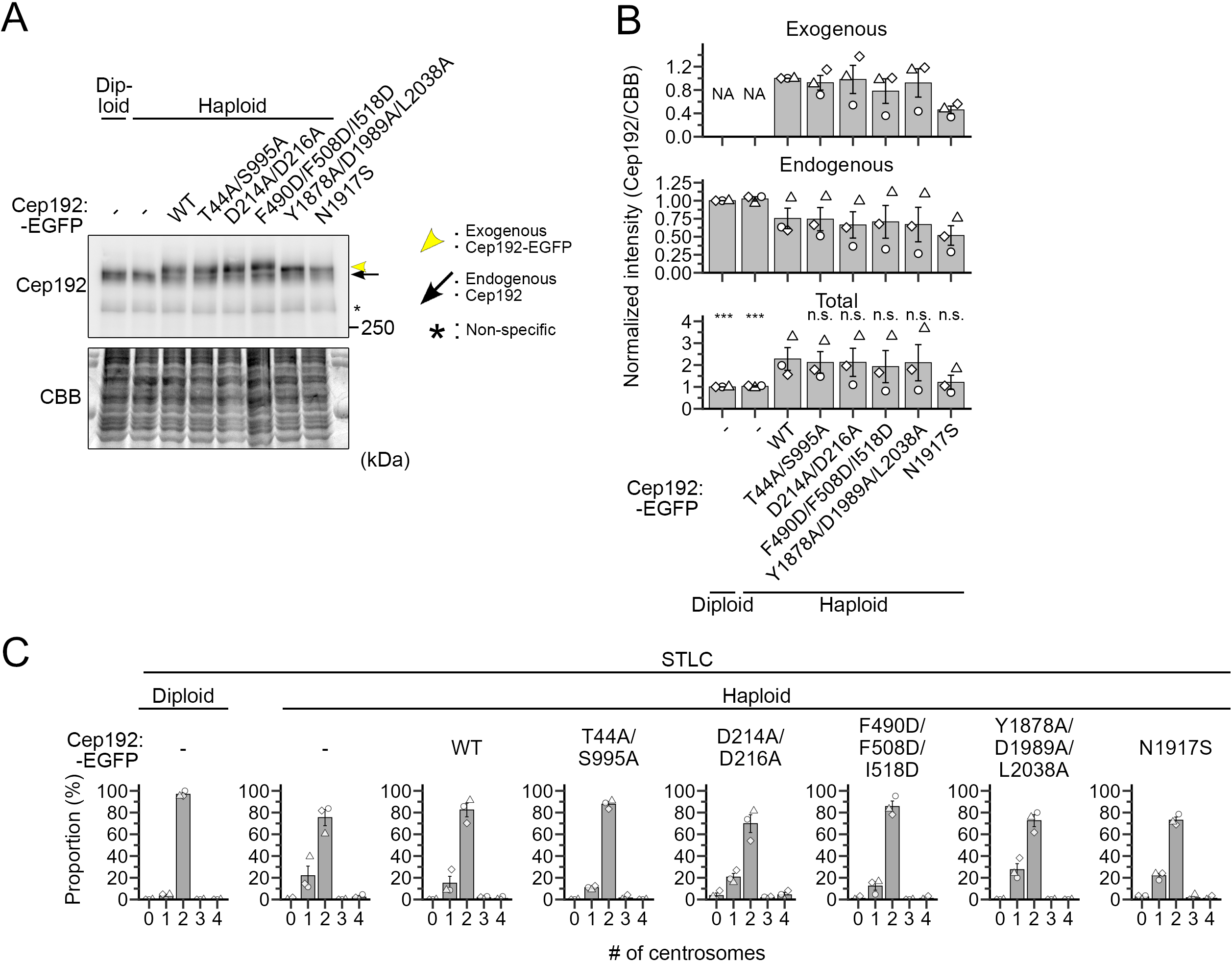
Exogenous expression of transgenic Cep192 mutants in haploids. **(A)** Immunoblotting of Cep192 in haploid or diploid cells with the indicated backgrounds. CBB-stained total protein was quantified as the loading control. **(B)** Quantification of the relative intensity of total Cep192 (i.e., summation of endogenous and exogenous transgenic Cep192 signals) in A. Protein loading differences were corrected based on CBB signals. Asterisks indicate statistically significant differences from haploid Cep192 (WT)-EGFP control (*** p<0.001, the Steel test). **(C)** Proportion of centrosome number in Fig. 5C. Mean ± SE of 3 independent experiments. At least 63 cells were analyzed for each condition.

**Fig. S5:**
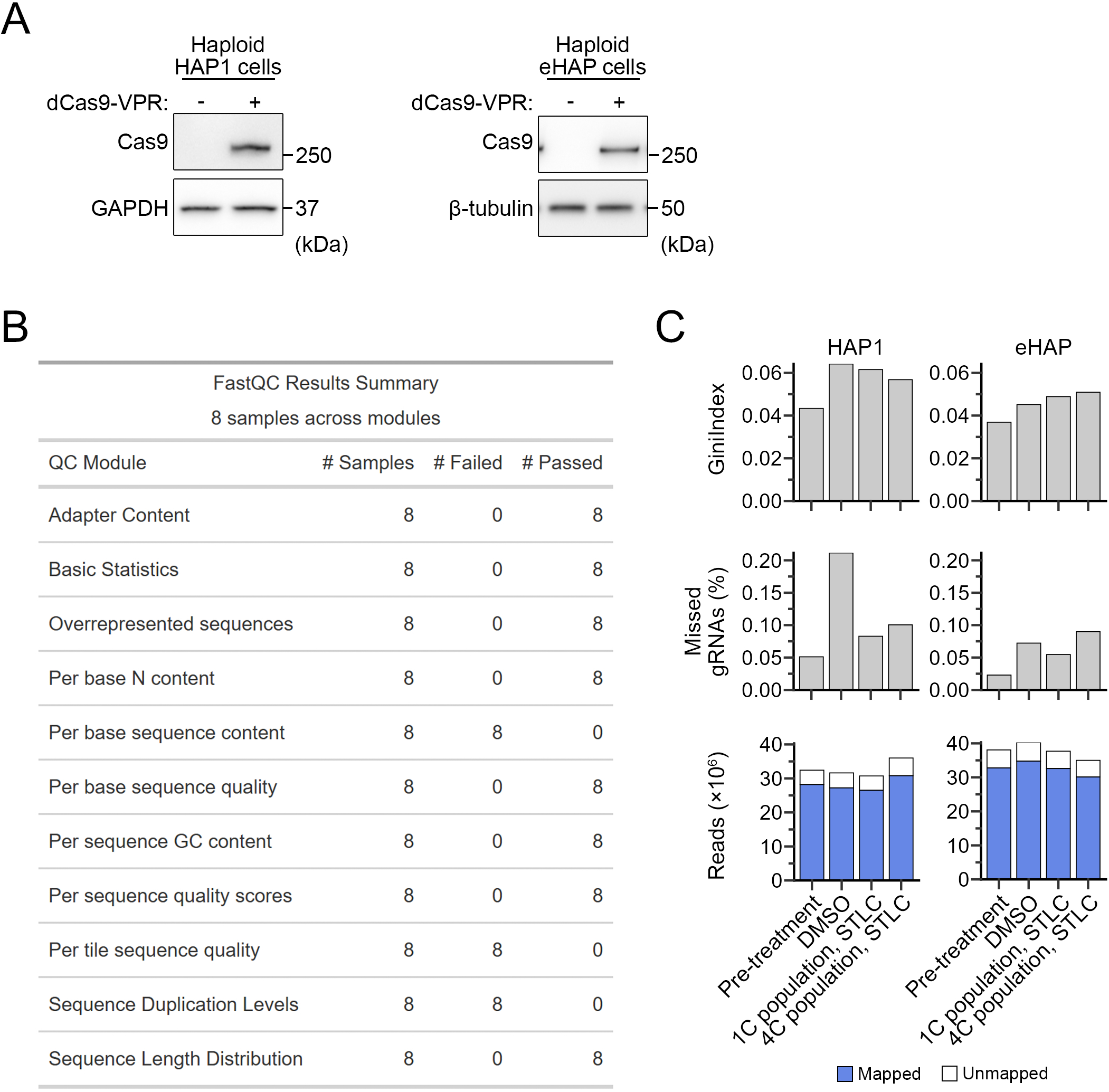
Quality check of the genome-wide CRISPRa screen. **(A)** Immunoblotting of dCas9-VPR using anti-Cas9 antibody in haploid HAP1 or eHAP cells with the indicated backgrounds. GAPDH or β-tubulin was detected as a loading control. **(B, C)** Summary of quality check for gRNA sequencing.

**Fig. S6:**
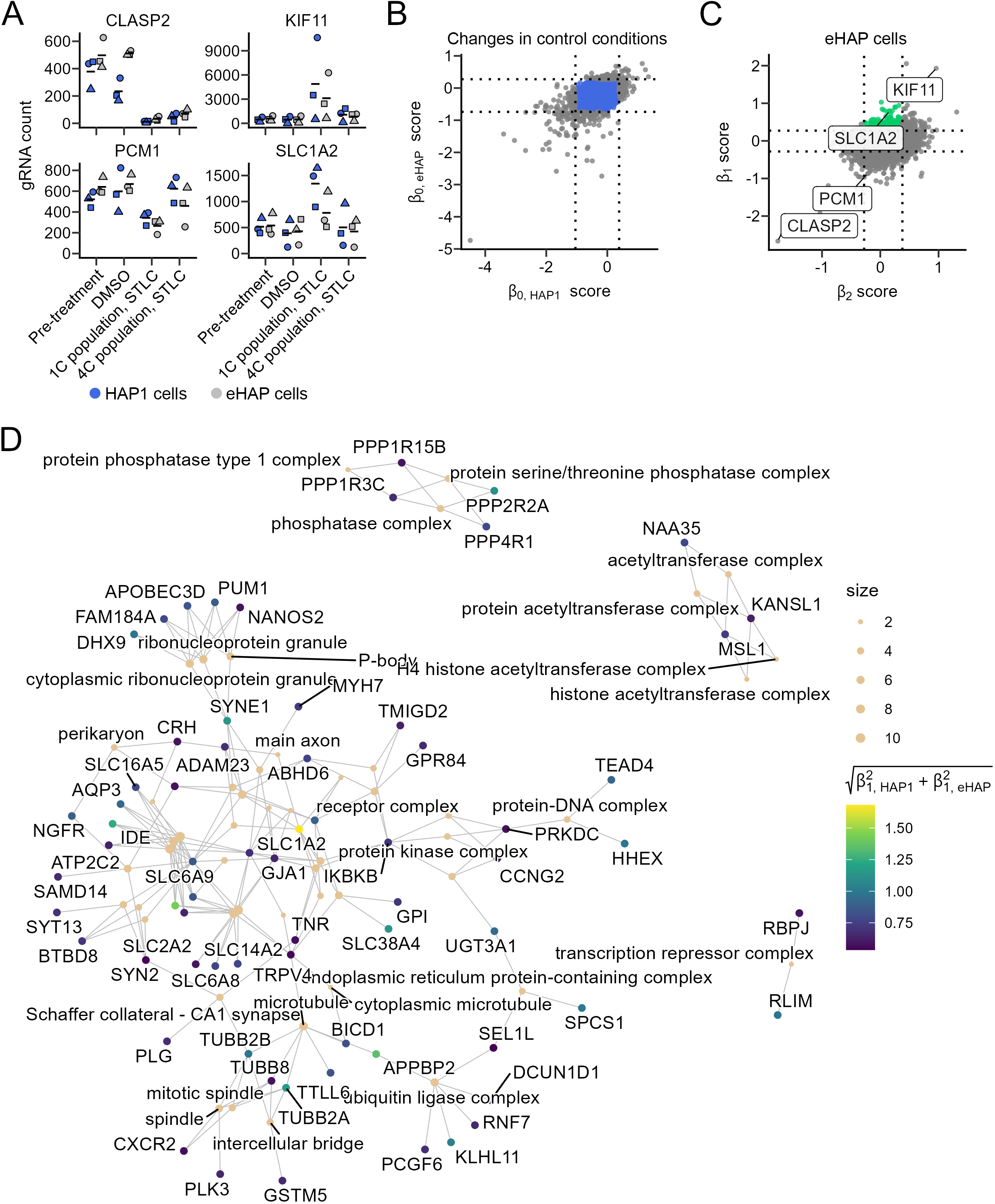
Result summary of the genome-wide CRISPRa screen. **(A)** Examples of gRNA count for several genes with distinct patterns of distribution in four different cell populations in the CRISPRa screen. **(B)** Graphical representation of the filtering out of genes with prominent fluctuations in their retention between pre-treated and vehicle control cell populations. Removed genes are indicated as grey dots. **(C)** A 2-D scatter plot of β_1_ vs β_2_ scores in eHAP cells. Hit genes selectively concentrated in the haploid G1 population under modest Eg5 inhibition were indicated in green. **(D)** GO terms of cellular component shared among at least 2 of 133 hit genes in Fig. 7C.

**Fig. S7:**
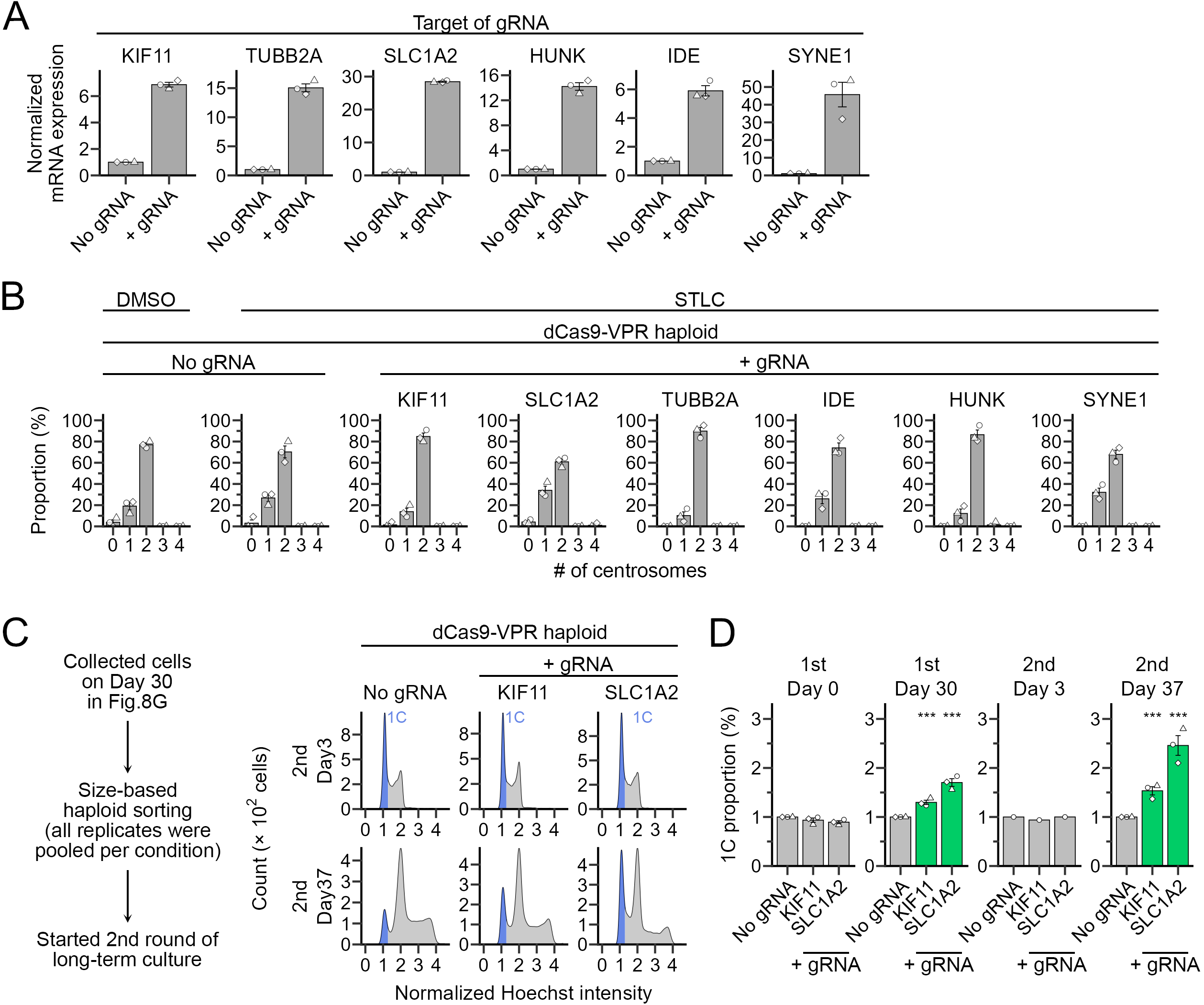
Identification of haploid-stabilizing genes. **(A)** Relative expression levels of gRNA target genes (normalized to that of β-actin) quantified by qPCR. Mean ± SE of 3 independent experiments. **(B)** Proportion of centrosome numbers in Fig. 8C. Mean ± SE of 3 independent experiments. At least 68 cells from 3 independent experiments were analyzed. **(C)** Flow cytometric analysis of DNA content in haploid dCas9-VPR cells expressing gRNA for the indicated genes during the second round of consecutive passages (the experimental flow is shown on the left). DNA was stained with Hoechst 33342. **(D)** Proportion of 1C populations in C. Mean ± SE of 3 independent experiments. Asterisks indicate statistically significant differences from the untransfected control (*** p<0.001, the Steel test). Identical data for the “1^st^ Day 30” sample are shown in Fig. 8H.

**Table S1:** gRNA count in the genome-wide CRISPRa screen

**Table S2:** β scores and pHaplo score for all genes analyzed in the genome-wide CRISPRa screen

**Table S3:** Cell lines established or used in this study

**Table S4:** Plasmids constructed or used in this study

**Table S5:** Primers used in this study

**Table S6:** Antibodies and compounds used in this study

**Supplementary Material S1: R script for the automated cell population quantification from flow cytometry data**

