## Supplementary Material S1 for "Cep192 insufficiency underlies haploid instability in human cells": SupplementalMaterial_S1_Yoshizawa_et_al_2026.Rmd

````
---
title: "R Notebook"
output: html_document
---

```{r import library and directory}
if (!require("pacman")) install.packages("pacman")

pacman::p_load(ggplot2, lemon, scales, tidyr, patchwork, data.table, readr, stringr, tcltk)

datadir <- tcltk::tk_choose.dir(caption = "Select the directory where the Rmd file is located.")
inputdir <- paste0(datadir, "/input/")
opdir <- paste0(datadir, "/output/")
```

```{r import data}
files <- list.files(path = inputdir)
csv.files <- grep("\\.csv$",files)

df <- data.frame(matrix(ncol = 14, nrow = 1))[numeric(0),] 

for(i in 1:length(files)){
  tmp <- read_csv(paste0(inputdir,"\\",files[i]), show_col_types = FALSE) %>% dplyr::mutate(LotNo = files[i])
  df <-rbind(df, tmp)
}

df_FCM <- df %>%
  dplyr::select(`FL7-H`, LotNo) %>% 
  dplyr::mutate(Cell = case_when(
    str_detect(LotNo, regex("mixed", ignore_case = TRUE)) ~ "Mixed_ploidy",
    str_detect(LotNo, regex("haploid", ignore_case = TRUE)) ~ "Haploid",
    str_detect(LotNo, regex("diploid", ignore_case = TRUE)) ~ "Diploid",
    TRUE ~ NA
  )) %>% 
  dplyr::select(-LotNo)

write.csv(df_FCM, file.path(datadir, "/df_FCM_Yoshizawa_et_al_biorxiv.csv"), row.names = FALSE)
df_FCM <- read_csv(paste0(datadir, "/df_FCM_Yoshizawa_et_al_biorxiv.csv"), show_col_types = FALSE)

```

```{r Definition of function}
local_maxima <- function(y, min_height_rel = 0.01, min_distance = 1, smooth_spar = 0.3) {
  if (smooth_spar > 0) {
    y <- smooth.spline(y, spar = smooth_spar)$y
  }
  
  dy <- diff(y)
  cand <- which(diff(sign(dy)) == -2) + 1
  global_max <- max(y)
  cand <- cand[y[cand] >= min_height_rel * global_max]
  
  if (length(cand) > 1 && min_distance > 1) {
    dist_matrix <- abs(outer(cand, cand, "-"))
    diag(dist_matrix) <- min_distance
    keep_idx <- which(colSums(dist_matrix < min_distance) <= 1)
    cand <- cand[keep_idx]
  }
  cand
}

select_closest_peak <- function(peaks_in_range, ctrl_peak) {
  if (length(peaks_in_range) == 0) {
    return(NA)
  }
  if (length(peaks_in_range) == 1) {
    return(peaks_in_range[1])
  }
  # Multiple peaks: return the one closest to control peak
  distances <- abs(peaks_in_range - ctrl_peak)
  return(peaks_in_range[which.min(distances)])
}
```

```{r Check 2C and 1C peaks in a control condition}
CtrlCell <- "Mixed_ploidy"

bw_parameter <- 0.8

dna_intensity <- df_FCM %>% dplyr::filter(Cell == CtrlCell) %>% pull(`FL7-H`)

dens_raw_ctrl <- density(dna_intensity, kernel = "gaussian", bw = "nrd0")

bw_rough  <- dens_raw_ctrl$bw * bw_parameter
dens_default  <- density(dna_intensity, kernel = "gaussian", bw = bw_rough)

peak_idx_default <- local_maxima(dens_default$y, min_height = 0.05 * max(dens_default$y))

df_dens_ctrl <- tibble(x = dens_default$x, y = dens_default$y)
df_peaks_ctrl <- tibble(x = dens_default$x[peak_idx_default], y = dens_default$y[peak_idx_default])
df_phase <- tibble(dna_intensity = dna_intensity)


p_hist <- ggplot() +
        geom_histogram(data = df_phase, aes(x = dna_intensity), binwidth = dens_default$bw/10, alpha = 0.8, position = "stack", linewidth = 0.01, fill = "darkgrey") +
        geom_line(data = df_dens_ctrl, aes(x = x, y = y * 2e4), linewidth = 0.4, alpha = 0.4, color = "black") +
        geom_point(data = df_peaks_ctrl, aes(x = x, y = y * 2e4), size = 1, alpha = 0.6, color = "black") +
        geom_label(data = df_peaks_ctrl, aes(label = round(x,1) , y = y * 2.5e4, x = x), vjust = -1, size = 5, fill = "white") +
        labs(x = "DNA intensity (a.u.)", y = "Count", title = paste0("Control plot")) +
        scale_x_continuous(limits = c(0, 700), breaks = seq(0, 700, 200)) +
        scale_y_continuous(expand = c(0,0)) +
        theme_bw()
p_hist
```

```{r Peak detection} 
#=====================
# USER INPUT
#=====================
Ctrl_2C_abs <- 302.3
Ctrl_1C_norm <- 161.9 / 302.3 * 2
sdv_default <- 0.15
bw_parameter <- 0.8
CtrlCellName <- "Mixed_ploidy"


#=====================

cat("Unique values:\n")
cat("Cells:", unique(df_FCM$Cell), "\n\n")

cells <- unique(df_FCM$Cell)

if (!dir.exists(opdir)) dir.create(opdir, recursive = TRUE)

df_dens_list <- list()
df_phase_list <- list()
df_peaks_list <- list()
count_idx <- 1


for (cell in cells) {

  dna_intensity_raw <- df_FCM %>% dplyr::filter(Cell == cell) %>% pull(`FL7-H`)
  
  if (length(dna_intensity_raw) == 0) {
    cat("Skipping:", cell, "(no data)\n")
    next}
  
  dens_raw <- density(dna_intensity_raw, kernel = "gaussian", bw = "nrd0")
  bw_rough  <- dens_raw$bw * bw_parameter
  dens_rough  <- density(dna_intensity_raw, kernel = "gaussian", bw = bw_rough)
  
  peak_idx_raw <- local_maxima(dens_rough$y, min_height = 0.05 * max(dens_rough$y))
  
  df_peaks <- tibble(x = dens_rough$x[peak_idx_raw], y = dens_rough$y[peak_idx_raw], bw = unique(dens_rough$bw))
  df_dens <- tibble(x = dens_rough$x, y = dens_rough$y, bw = unique(dens_rough$bw))
  
  peak_2c_candidates <- df_peaks$x[df_peaks$x >= Ctrl_2C_abs * 0.85 & df_peaks$x <= Ctrl_2C_abs * 1.15]
  peak_2c_idx_raw <- select_closest_peak(peak_2c_candidates, Ctrl_2C_abs)
  peak_2c_idx <- peak_2c_idx_raw / (peak_2c_idx_raw / 2)
  
  df_peaks$x_norm <- df_peaks$x / (peak_2c_idx_raw / 2)
  df_dens$dna_intensity_norm <- df_dens$x / (peak_2c_idx_raw / 2)
  
  peak_1c_candidates <- df_peaks$x_norm[df_peaks$x_norm >=  0.85 & df_peaks$x_norm <= 1.15]
  peak_1c_idx <- select_closest_peak(peak_1c_candidates, Ctrl_1C_norm)
  
  
  mu  <- c(OneC = peak_1c_idx, TwoC = peak_2c_idx)
  sdv <- c(OneC = sdv_default * peak_1c_idx / peak_2c_idx, 
           TwoC = sdv_default)
  
  ci <- data.frame(
    Phase = c("1C", "2C"), 
    mu = mu, 
    sd = sdv,
    lower = qnorm(0.01, mean = mu, sd = sdv),
    upper = qnorm(0.99, mean = mu, sd = sdv)
  )
  
  L1 <- ci$lower[1]; U1 <- ci$upper[1]
  L2 <- ci$lower[2]; U2 <- ci$upper[2]
  
  if (length(peak_2c_idx) == 0 || is.na(peak_2c_idx)) {
    cat("Warning:", rep, "-", treat, "-", cell, "(no 2C peak found in expected range)\n")
    next
  }
  
  df_dens <- df_dens %>% 
    dplyr::mutate(area_group = if (is.na(peak_1c_idx)) {
      case_when(dna_intensity_norm <= U2  ~ "2C",
                TRUE ~ "Other")
    } else {
      case_when(dna_intensity_norm <= U1 ~ "1C",
                TRUE ~ "Other")
    })
  
  df_phase <- tibble(dna_intensity = dna_intensity_raw) %>%
    dplyr::mutate(
      dna_intensity_norm = dna_intensity_raw / (peak_2c_idx_raw / 2), 
      area_group = if (is.na(peak_1c_idx)) {
      case_when(dna_intensity_norm <= U2  ~ "2C",
                TRUE ~ "Other")
    } else {
      case_when(dna_intensity_norm <= U1 ~ "1C",
                TRUE ~ "Other")
    })
  
  cat("Processed:", cell, "\n")
  if (is.na(peak_1c_idx)) {
    cat("  -> Using NO-1C mode (2C peak detected at", round(peak_2c_idx, 2), ")\n")
  } else {
    cat("  -> Using FULL mode (1C peak detected at", round(peak_1c_idx, 2), ")\n")
  }
  
  df_dens_list[[count_idx]] <- df_dens %>% dplyr::mutate(Cell = cell) 
  df_phase_list[[count_idx]] <- df_phase %>% dplyr::mutate(Cell = cell) 
  df_peaks_list[[count_idx]] <- df_peaks %>% dplyr::mutate(Cell = cell) 
  
  count_idx <- count_idx + 1
}
      
df_dens_all <- bind_rows(df_dens_list)
df_phase_all <- bind_rows(df_phase_list)
df_peaks_all <- bind_rows(df_peaks_list)

write.csv(df_dens_all, file.path(paste0(opdir, "demo_df_dens_all.csv")), row.names = FALSE)
write.csv(df_phase_all, file.path(paste0(opdir, "demo_df_phase_all.csv")), row.names = FALSE)
write.csv(df_peaks_all, file.path(paste0(opdir, "demo_df_peaks_all.csv")), row.names = FALSE)
```

```{r plot}
xorder <- c("Haploid", "Mixed_ploidy", "Diploid")

p_hist <- 
  ggplot(df_FCM, aes(x = `FL7-H`)) +
  facet_wrap2(.~factor(Cell, levels = xorder), scales = "free_y", axes = "all") +
  geom_histogram(binwidth = 2, alpha = 0.8, position = "stack", linewidth = 0.01, fill = "darkgrey") +
  labs(x = "Absolute DNA intensity (a.u.)", y = "Count", title = "1. Input") +
  scale_x_continuous(limits = c(0, 700), breaks = seq(0, 700, 200)) +
  scale_y_continuous(expand = expansion(mult = c(0, 0.1))) +
  theme_bw()

p_hist_norm <- 
  ggplot(df_phase_all, aes(x = dna_intensity_norm)) +
  facet_wrap2(.~factor(Cell, levels = xorder), scales = "free_y", axes = "all") +
  geom_histogram(binwidth = dens_rough$bw/400, alpha = 0.8, position = "stack", linewidth = 0.01, fill = "darkgrey") +
  geom_point(data = df_peaks_all, aes(x = x_norm, y = y * 5e4), size = 1, alpha = 0.6, color = "black") +
  labs(x = "Normalized DNA intensity (a.u.)", y = "Count", title = "2. Peak detection using the algorithm") +
  scale_x_continuous(limits = c(0, 4.5), breaks = seq(0, 4, 1)) +
  scale_y_continuous(expand = expansion(mult = c(0, 0.1))) +
  theme_bw()

p_dens <- 
  ggplot(df_dens_all, aes(x = dna_intensity_norm)) +
  facet_wrap2(.~factor(Cell, levels = xorder), scales = "free_y", axes = "all") +
  geom_line(aes(y = y * 10e4), linewidth = 0.2, alpha = 1, color = "black") +
  geom_area(aes(y = y * 10e4, fill = area_group), alpha = 0.6) +
  scale_fill_manual(values = c("royalblue","royalblue", "grey")) +
  labs(x = "Normalized DNA intensity (a.u.)", y = "Count", title = "3. Output") +
  scale_x_continuous(limits = c(0, 4.5), breaks = seq(0, 4, 1)) +
  scale_y_continuous(expand = expansion(mult = c(0, 0.1))) +
  theme_bw() +
  theme(legend.position = "none")

p <- p_hist / p_hist_norm / p_dens
p

ggsave(file = "Peak_detection_demo_Yoshizawa_et_al_2026_biorxiv.png", plot = p, device = "png", path = opdir, width = 5, height = 6, dpi = 300, limitsize=FALSE)
```
````
